# Spatial ploidy inference using quantitative imaging

**DOI:** 10.1101/2025.03.11.642217

**Authors:** Nicholas J. Russell, Paulo B. Belato, Lilijana Sarabia Oliver, Archan Chakraborty, Adrienne H. K. Roeder, Donald T. Fox, Pau Formosa-Jordan

## Abstract

Polyploidy (whole-genome multiplication) is a common yet under-surveyed property of tissues across multicellular organisms. Polyploidy plays a critical role during tissue development, following acute stress, and during disease progression. Common methods to reveal polyploidy involve either destroying tissue architecture by cell isolation or by tedious identification of individual nuclei in intact tissue. Therefore, there is a critical need for rapid and high-throughput ploidy quantification using images of nuclei in intact tissues. Here, we present iSPy (<u>I</u>nferring <u>S</u>patial <u>P</u>loid<u>y</u>), a new unsupervised learning pipeline that is designed to create a spatial map of nuclear ploidy across a tissue of interest. We demonstrate the use of iSPy in Arabidopsis, Drosophila, and human tissue. iSPy can be adapted for a variety of tissue preparations, including whole mount and sectioned. This high-throughput pipeline will facilitate rapid and sensitive identification of nuclear ploidy in diverse biological contexts and organisms.

## Introduction

Polyploidy, whereby cells have more than two homologous copies of their chromosomes, can either occur in every somatic cell of an organism (i.e., organismal polyploidy) or a subset of cells (i.e., endoploidy or endopolyploidy)^1^. Endopolyploidy commonly arises through repeated cell cycles where DNA replication occurs without completed cell division (referred to by many names such as endoreplication) or by cell-to-cell fusion^2^. Endopolyploid tissues are widespread among eukaryotic organisms and can be spatially patterned in both plants^3–5^ and animals^6–8^. Endopolyploidy can occur during organ formation^4,7,9^ or it can be promoted through external stimuli, for example, to regenerate tissue after injuries or aid in immune response^1,10,11^. New examples of endopolyploidy are continually being identified, revealing that polyploidy is a common intrinsic property of most tissue types. However, ectopic occurrences of endopolyploidy can elevate the risks of genome instability and diseases such as cancer^2,12–14^. Therefore, identifying abnormal endopolyploid cells in tissue biopsy samples is often crucial.

Given the ever-increasing appreciation of the importance of endopolyploidy in tissue biology, powerful methods should be available to accurately quantify nuclear ploidy in a tissue of interest at a given time and position. Preferably, this quantification should occur while the tissue or organism is still intact, to reveal the position of the polyploid cells. Flow cytometry is a frequently used approach to measure the ploidy distribution in tissues in both plants^15–18^ and animals^19,20^. However, this method is invasive and the sampled tissue is destroyed in the process, making it very difficult to recover positional information. Computational techniques have been developed to infer ploidy from sequencing data, but this also removes spatial information and is a relatively low-throughput method^21–24^.

Alternatively, high-resolution microscopy followed by single-cell analysis can give accurate measurements of ploidy while retaining spatial context^25–27^. However, manual analyses of nuclear ploidy from microscopy imaging data are tedious and low-throughput, because each nucleus is measured individually. Other common methods of labeling DNA content while using fluorescence microscopy techniques include fluorescence *in situ* hybridization^28^ or DNA stains such as propidium iodide (PI), Hoechst, or 4’,6-diamidino-2-phenylindole (DAPI)^9,17,29–31^. Pipelines have also been created to streamline the quantification of ploidy from microscopy images, some using deep learning methods^9,17,32^. However, these methods remain relatively low throughput due to the large amount of background noise from PI– and DAPI-staining, necessitating laborious manual quantification of nuclear fluorescence.

In this paper, we introduce iSPy (Inferring Spatial Ploidy), a high-throughput pipeline to quantify the ploidy of nuclei while retaining spatial information. We use confocal imaging techniques that preserve tissues, and an unsupervised Gaussian mixture model to predict the ploidy of all nuclei and to produce a ploidy spatial map. We demonstrate the efficacy of this technique using three distinct model systems. First, we highlight the utility of iSPy to pinpoint developmentally programmed endopolyploidy in plant tissue: the cotyledons of *Arabidopsis thaliana* (hereafter Arabidopsis). We verify that iSPy-derived data from this tissue closely matches data from flow cytometry. Second, we show that iSPy can track regeneration-induced endopolyploidy in a physically compressed sample from the *Drosophila melanogaster* (hereafter Drosophila) hindgut pylorus. Third, we highlight the ability of iSPy to track ploidy differences in physically sectioned samples from human organ donor hearts. Our data reveal the broad applicability of iSPy and its ability to identify complex spatial positioning of cells with different ploidies. This allows researchers across several fields to accurately and rapidly quantify ploidy across tissues in diverse organisms and enhances the ability to identify and analyze endopolyploid cells.

## Results

### A machine learning-based image analysis method to determine nuclear ploidy

The starting point for the iSPy pipeline is a tissue preparation containing a fluorescence marker or dye that correlates with DNA content (Fig. 1A). As discussed in later sections, our technique applies to a wide variety of tissue preparations from diverse organisms. From each tissue preparation, we acquired a three-dimensional confocal image with appropriate nuclear markers. We segmented the nuclei from either the raw three-dimensional confocal image or from the two-dimensional sum-projected image to obtain key nuclear features using the high-throughput image segmentation software ilastik, interactively segmenting nuclei using supervised classification and thresholding (see Methods and subsequent subsections for more specifics)^33^. For a system such as Arabidopsis, three-dimensional segmentation is preferred due to the presence of multiple cell types at distinct depths within the tissue. However, for Drosophila and human cardiomyocytes, two-dimensional segmentation of the sum-projected images is feasible due to sample preparation techniques. After nuclear segmentation was completed, we manually labeled relevant objects, such as known cell types and erroneous nuclear segmentations, using ilastik’s Object Classification tool. Lastly, we extracted nuclear features such as the intensity of the corresponding nuclear markers and the nuclear volume (Fig. 1B).

**Figure 1:**
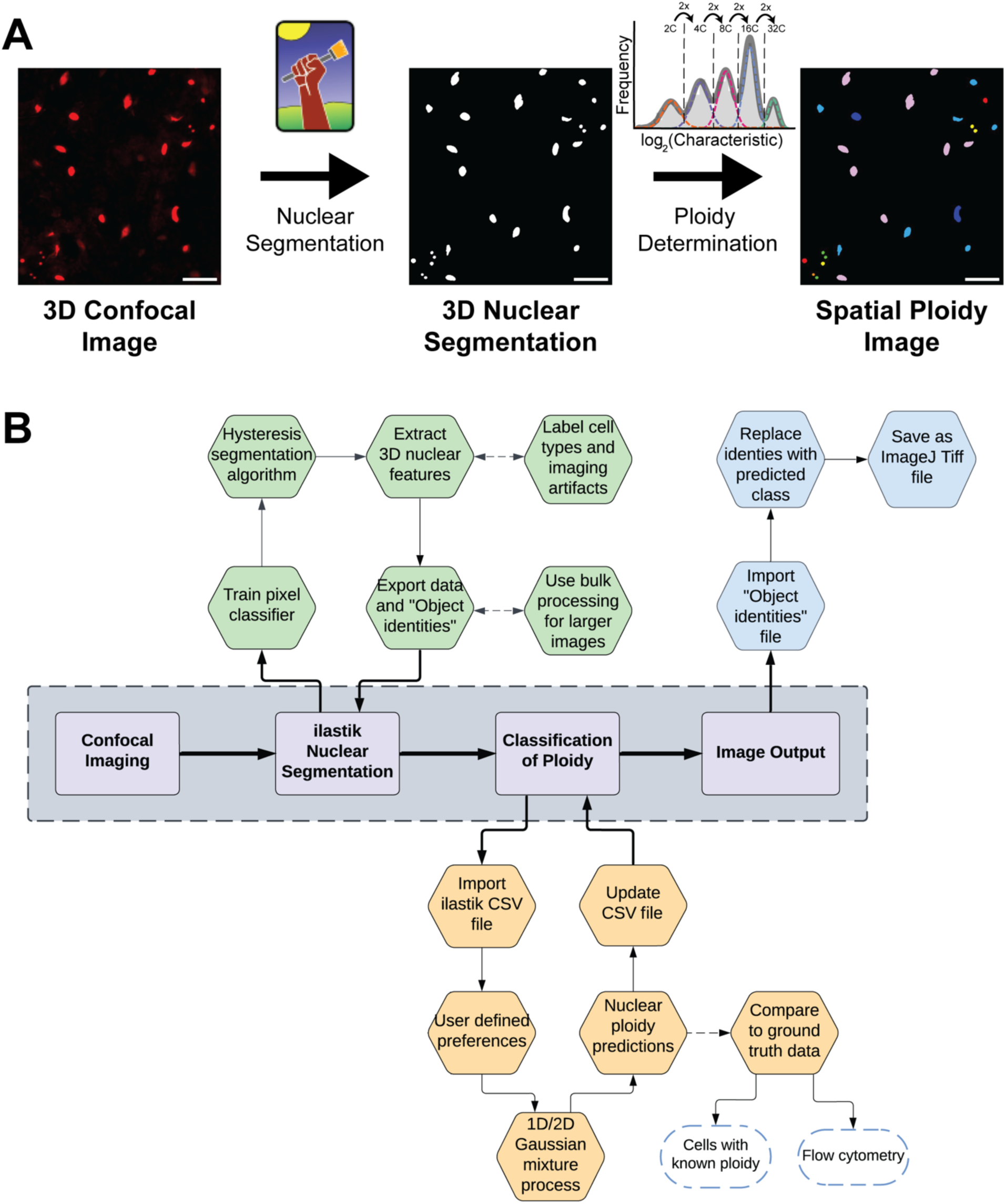
An image-based pipeline to determine spatial ploidy. **(A)** The iSPy pipeline at a glance. Confocal imaging is performed with nuclear reporters and stains, nuclear segmentation with ilastik is conducted, and a Gaussian mixture model (Gaussian mixture) is used to classify the ploidy of each nucleus with easy-to-use software. Colors in the right panel represent different ploidy levels. Scalebars = 25 µm. **(B)** A more detailed overview of the iSPy procedure. See the Results and Methods sections for further details. Dashed arrows indicate optional processes and are not essential to use iSPy for analyses. See Fig. S1 for a more detailed procedure for the Gaussian mixture process.

To identify clusters of nuclei with equal ploidy, we utilized the unsupervised Gaussian mixture model^34^. This method assumes that the features we are interested in come from the combination of several normally distributed populations, and these populations correspond to the different ploidy classes (1C corresponding to the haploid genome, 2C corresponding to a diploid genome, 4C, 8C, etc.). One of the key features we were interested in was the total intensity of the nuclear markers. Because we expect the nuclear marker signal to double as genome copy number doubles (e.g., if 2C nuclei have a mean total intensity of 2^k^, we expect 4C nuclei to have a mean of 2^k+1^, 8C with 2^k+2^, etc.), we scaled all of our data logarithmically with base 2, similar to other ploidy quantification pipelines^22,23,30^. We then input the data into our Python software and output the best-fit model by varying the parameters of our model. To identify the Gaussian mixture model that provided the best fit for our data, we implemented well-established minimization metrics such as the Akaike information criterion (AIC) and the Bayesian information criterion (BIC) (see Methods)^34,35^. We then classified each nucleus to one of the clusters by using its corresponding log-likelihood probabilities (see Methods).

Thus, our pipeline classifies and reports the ploidy of each nucleus within an input image and generates a two-or three-dimensional segmented image of the resulting classification, as well as data files suitable for further analyses of the spatial distribution of ploidies (Fig. 1). In subsequent sections, we highlight three diverse examples of tissues where we applied our method to quantify tissue ploidy.

### iSPy facilitates ploidy determination in plant tissues expressing fluorescent reporters

In Arabidopsis, the ploidy level in cells of sepals, leaves, and cotyledons varies^36–39^. Endopolyploidy is initiated at different time points throughout development, resulting in a heterogeneous ploidy distribution throughout the tissue, with cells varying from 2C to 64C^36,40^. Previously, classification of cellular ploidy was difficult without performing flow cytometry.

Recent work has demonstrated general correlations between nuclear sizes, cell sizes, and ploidy, but there are exceptions in both sepals and leaves^36,37^. Many studies have used PI or DAPI staining to assist in calculating the nuclear ploidy of fixed tissues, although there is no standardized method to quantify ploidy from these types of images^9,37^. Moreover, live, undamaged cells are often impermeable to PI and other dyes, meaning these stains cannot be used for single-nuclei tracking of ploidy in time-lapse microscopy of living tissues. To accurately determine the ploidy of living nuclei throughout tissue development, it is necessary to develop new techniques that do not require tissue destruction or fixation. Recently, it has been shown that the presence of histone markers in yeast correlates with genome content, but whether the same holds in plants – and in particular Arabidopsis – has not been explored in depth^41^.

To this end, we harvested 14-day-old cotyledons of plants expressing the fluorescently tagged nuclear histone marker (*p35S::H2B-RFP1*; Cauliflower Mosaic virus promoter 35S driving expression of a Histone 2B red fluorescent protein fusion) to quantify ploidy (Fig. 2A; Methods). We performed confocal microscopy, segmented the three-dimensional nuclei using ilastik, and hand-selected stomatal guard cell nuclei of known 2C ploidy as a standard (Fig. 2A; Methods). We plotted frequency histograms of the nuclear volume and the total nuclear signal intensity of each nucleus, and we observed five distinct peaks corresponding to 2C, 4C, 8C, 16C, and 32C nuclei (Fig. S2A). These peaks or clusters were also apparent when considering both volume and intensity simultaneously. The stomatal guard cell nuclei, an internal standard known to be 2C, formed a cluster along with other epidermal nuclei, so we assigned this peak as 2C. However, the majority of nuclei were endoreduplicated and were 4C and above. Note that not all fluorescent nuclear markers produce evident peaks, likely due to differences in the expression level of the promoter in different cells. For example, the epidermal-specific nuclear marker *pML1::H2B-mTFP* (*Arabidopsis thaliana MERISTEM LAYER1* promoter driving expression of Histone 2B fused with teal fluorescent protein) does not have evident ploidy clusters (Fig. S2B–C). The only cluster evident contained the stomatal guard cell nuclei, which had a much lower total fluorescent intensity and volume than other epidermal nuclei, correlating with the known low level of expression of the pML1 promoter in guard cells (Fig. S2B–C, arrow; Methods)^42^. Thus, we continued to use *p35S::H2B-RFP1* in our ploidy analysis.

**Figure 2:**
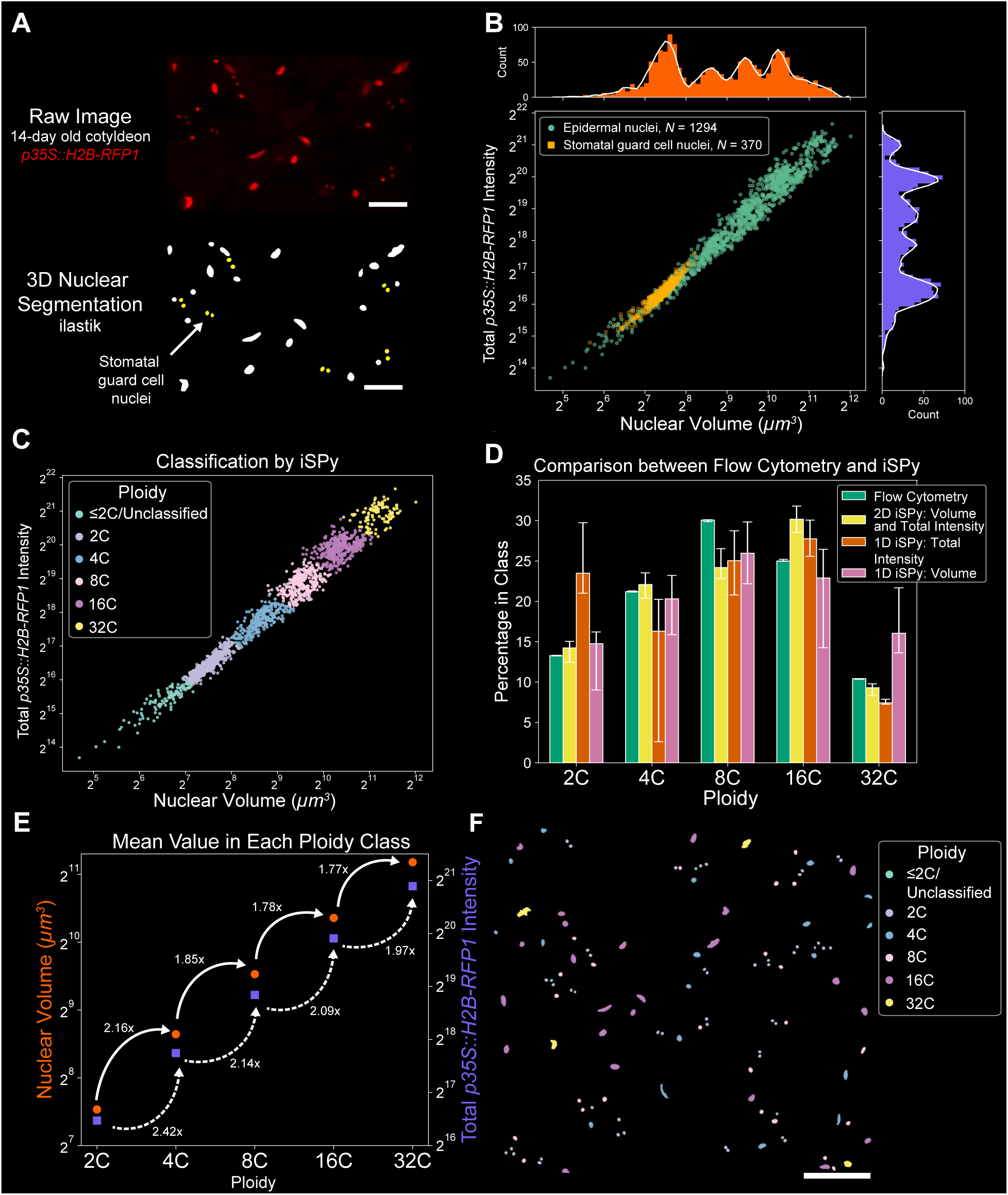
iSPy shows that the histone marker accurately quantifies ploidy in Arabidopsis cotyledons. **(A)** Above: representative sum-projected confocal image of 14-day-old Arabidopsis cotyledons with the nuclear marker *p35S::H2B-RFP1*. Below: 3D nuclear segmentation of the above image performed in ilastik. Yellow nuclei signify hand-selected stomatal guard cell nuclei (see Methods). Scale bar = 25 µm. **(B)** Scatter plot of the nuclear volume and total *p35S::H2B-RFP1* intensity of the ilastik-segmented epidermal nuclei (green) and stomatal guard cell nuclei (yellow) including corresponding histograms of the total *p35S::H2B-RFP1* intensity (purple) and nuclear volume (orange) with a smoothed Savitzky–Golay filter (white line, only for illustrative purposes). **(C)** iSPy prediction using the 2D Gaussian mixture with six components and spherical covariance matrix (see Fig. S5A–B for additional information). **(D)** Comparison of the percentage of epidermal nuclei predicted in each ploidy class between flow cytometry (green, Fig. S2D–G), the 2D Gaussian mixture with nuclear volume and total intensity (yellow, Fig. 2C, S5A–B), the 1D Gaussian mixture with only total intensity (red, Fig. S4A–C), and the 1D Gaussian mixture with only nuclear volume (pink, Fig. S4D–F). See Supplemental Data Set 1 for exact values. Uncertainty bars represent nuclei that may be classified incorrectly (log-likelihood probability less than 0.8) and nuclei that could be classified in another component (log-likelihood probability greater than 0.2) (see Methods). **(E)** The mean of the total *p35S::H2B-RFP1* intensity (purple squares, left axis) and nuclear volume (orange circles, right axis) with the fold increase to the next ploidy class using 2D iSPy in (C). **(F)** Segmented nuclear image colored with the ploidy distribution predicted from 2D iSPy in (C) across an abaxial side of a cotyledon from the same sample as in the inset shown in A.

We compared our iSPy results with those from flow cytometry performed on 14-day-old cotyledons (Fig. S2D–F). For flow cytometry, we utilized the epidermal-specific marker *pML1::H2B-mTFP* to distinguish epidermal from non-epidermal nuclei while simultaneously using PI staining to quantify DNA content. We observed a phenomenon similar to that in the confocal imaging data, with a cluster of stomatal guard cell nuclei showing low *pML1::H2B-mTFP* fluorescent intensity (Fig. S2C,F, arrows). Due to low *pML1::H2B-mTFP* intensity, it is possible that not all stomatal guard cell nuclei could be distinguished from non-epidermal nuclei. Therefore, to properly compare the flow cytometry data with our confocal imaging, we removed stomatal guard cell nuclei from flow cytometry and the sub-epidermal cells from our confocal imaging. For the flow cytometry analysis, we filtered the stomatal guard cell nuclei by setting a threshold on *pML1::H2B-mTFP* fluorescent intensity (Fig. S2F, dashed line; Methods). We obtained the ploidy distribution from flow cytometry by performing a Gaussian mixture using the total intensity of the PI staining in each nucleus (Fig. S2G; Methods). For the confocal imaging, we observed that sub-epidermal nuclei have a lower variance of intensity than epidermal nuclei, so we removed sub-epidermal nuclei by setting a threshold on the variance of intensity (Fig. S3; Methods). When considering only epidermal nuclei, the peaks from the frequency histograms of nuclear volume and the total nuclear signal intensity were more evident and the clusters when considering both nuclear features were clearer (Fig. 2B).

To cluster the ploidy of epidermal nuclei from our confocal images, we compared the performance of three Gaussian mixtures: a 1D Gaussian mixture using the total *p35S::H2B-RFP1* intensity (Fig. S4A–C), a 1D Gaussian mixture using the nuclear volume (Fig. S4D–F), and a 2D Gaussian mixture using both of these features (Figs. 2C and S5). Six components were optimal for all three Gaussian mixtures, but they varied in their classification due to the components in which the 2C stomatal guard cell nuclei were classified. For the 1D Gaussian mixture on total intensity, the two components with the lowest mean corresponded to 2C nuclei, and the remaining four peaks corresponded to 4C, 8C, 16C, and 32C (Fig. S4B–C). For the other two Gaussian mixtures, the component with the lowest mean corresponded to ≤2C/Unclassified nuclei which were subsequently filtered out, and the other five peaks corresponded to 2C, 4C, 8C, 16C, and 32C (Figs. 2C, S4E–F).

The ploidy distribution of epidermal cell nuclei using the Gaussian mixture models was comparable to the ploidy distribution obtained from the flow cytometry data (Figs. 2D, S5C–F). The 1D Gaussian mixtures, using either total nuclear signal or nuclear volume only, performed well for larger or smaller ploidies, respectively. However, using both of these features simultaneously in the 2D Gaussian mixture allowed for a more accurate classification across all ploidy levels, as shown by greater correspondence to flow cytometry results (Cramér–von Mises criterion against flow cytometry distribution: total signal, *T* = 1.642, *p* = 8.209×10^-5^; nuclear volume, *T* = 0.566, *p* = 0.0272; both features, *T* = 0.336, *p* = 0.10684; Figs. 2D, S5F). When using the 2D Gaussian mixture model, it was calculated that the total intensity of *p35S::H2B-RFP1* increased 2–2.4-fold as ploidy doubled, whereas the nuclear volume increased by 1.8–2.1-fold (Fig. 2E). This estimation is consistent with previous experimental and image processing work in the sepal^37^. After classification of the ploidy of each nucleus, iSPy outputs a maximal projection image showing the spatial arrangement of nuclear ploidy in the cotyledon epidermis (Fig. 2F). We observed several clusters of 4C, 8C, and 16C nuclei, while 32C nuclei were more isolated across the tissue and more likely to be farther away from stomata. Therefore, we found that the iSPy method using both total intensity and nuclear volume accurately predicts nuclear ploidy and offers a non-invasive technique to assess and visualize ploidy in leaves.

### iSPy accurately identifies injury severity-dependent endopolyploidy in the regenerating Drosophila pylorus

In certain animal tissues, regeneration after injuries can occur through the induction of endopolyploid cells^1,6,10^. Following an acute apoptotic injury in Drosophila, the surviving cells of the naturally diploid adult hindgut pyloric epithelium (hereafter, pylorus) endoreduplicate to restore tissue-wide DNA content to pre-injury levels^6,43,44^. Previously, we found that the degree of ploidy is tuned to the degree of injury in the pylorus, and more specifically, the ploidy of uninjured pyloric nuclei centers around 2C, mildly injured and fully regenerated pyloric nuclei centers around 4C, and severely injured pyloric nuclei are usually 8-16C^43^. Regardless of the degree of injury, the final regenerated tissue-wide ploidy is similar, thus showing that endoreduplication is coordinated with the degree of cell loss.

Quantification of ploidy in these experiments has previously been performed using an established protocol, which has proven to accurately determine ploidy in various Drosophila tissues^7,26,43,45,46^. Briefly, this protocol involves physically compressing (squashing) the pylorus, reducing the thickness and the variation in distance from the imaging objective to individual nuclei, and providing a more accurate ploidy quantification. Using this validated method, we previously detailed an accurate protocol whereby the ploidy of individual nuclei is defined and measured in Fiji^26^. This protocol provided spatial ploidy information; however, the manual segmentation of nuclei was labor intensive, thus prohibiting high-throughput analysis.

To determine whether iSPy could accurately produce spatial ploidy maps of regenerating Drosophila pylori, we used an established genetic cell ablation protocol to injure the pylorus and analyze tissue ploidy^26,43,44,46–48^. We examined three different conditions: uninjured pylori, mildly injured pylori (24 h at 29**°**C), and severely injured pylori (48 h at 29**°**C) (see Methods). As performed previously, we used an internal control, haploid (1C) spermatids were placed on the same slide (Fig. S6). We performed a sum projection of all images, segmented the projected two-dimensional nuclei in ilastik, and selected the pyloric (Fig. 3A) and spermatid (Fig. S6A) nuclei that were fully complete (see Methods). After extracting relevant nuclear information such as total Hoechst intensity and its projected area, we normalized the total Hoechst intensity of the pyloric nuclei to the median intensity of the haploid spermatids from the same experiment to account for variability between experiments as we have done previously (Fig. S6B, Methods).

**Figure 3:**
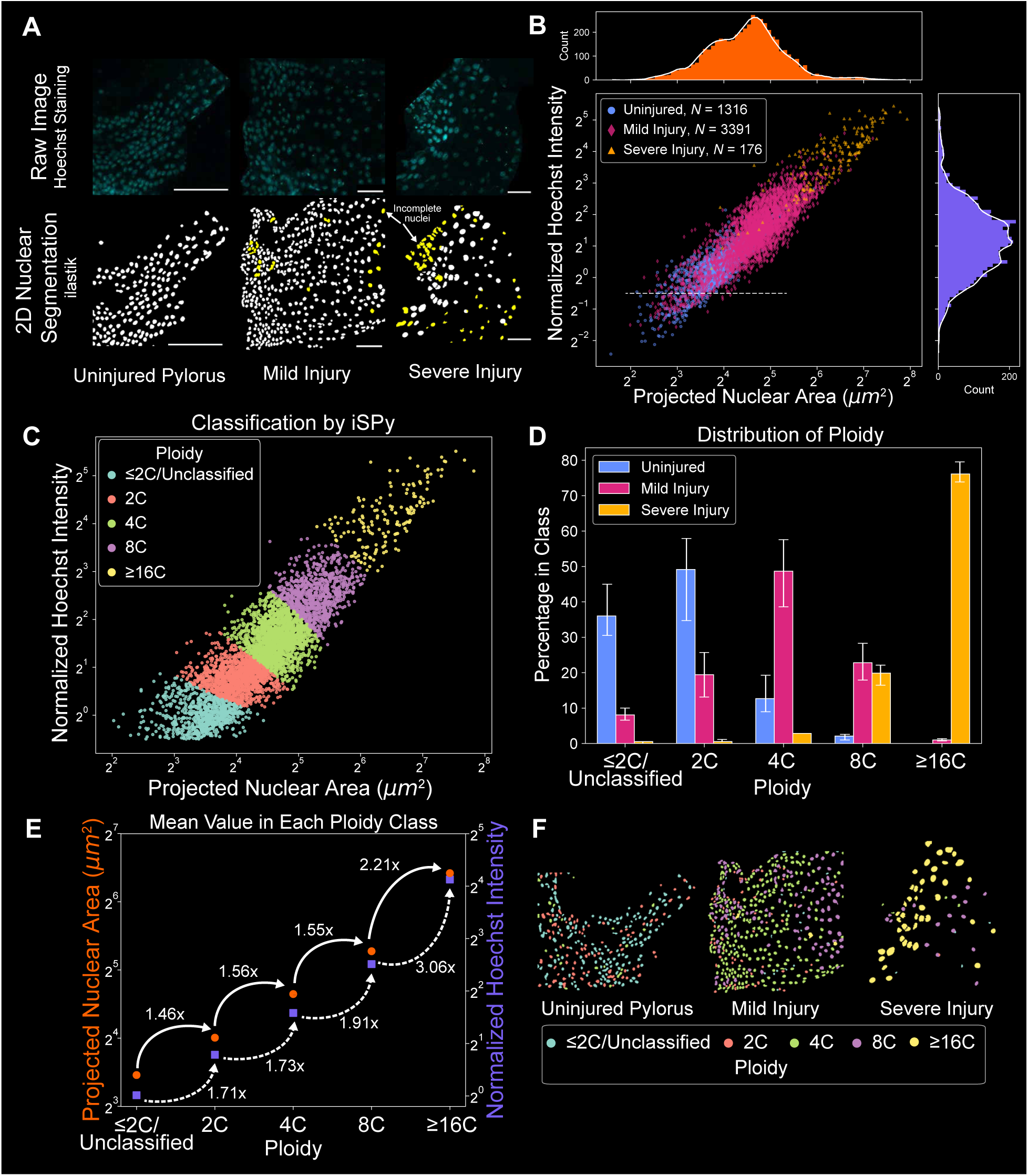
iSPy correctly predicts that Drosophila pyloric cells endoreduplicate in a severity-dependent manner. **(A)** Above: representative sum-projections of confocal images of pyloric nuclei stained with Hoechst (blue, see Methods) from an uninjured, mildly injured, and severely injured pylorus (left to right). Below: segmented nuclei using ilastik. Yellow segmented objects denote incomplete nuclei. Note that this segmentation was performed with the sum projection in two dimensions. The anterior and posterior sides of the tissue axis are on the left and right sides in each image, respectively. Scale bars: 50 µm. **(B)** Scatter plot of the projected nuclear area and normalized Hoechst intensity of the segmented nuclei (uninjured, blue circles; mild injury, red diamonds; severe injury, yellow squares) including corresponding histograms of the normalized Hoechst intensity (purple) and projected nuclear area (orange) with a smoothed Savitzky–Golay filter (white line, only for illustrative purposes). Cells that had a normalized Hoechst intensity below 2^-0.5^ were filtered out (dashed white line, see Methods). **(C)** iSPy ploidy prediction using the 2D Gaussian mixture with five components and diagonal covariance matrix (see Fig. S8A for additional information). Note the cut-off used from (B) at 2^-0.5^. **(D)** The proportion of pyloric cells in each ploidy class predicted from the 2D Gaussian mixture in (C) by severity of injury (uninjured, blue; mild injury, red; severe injury, yellow). Uncertainty bars represent nuclei that may be classified incorrectly (log-likelihood probability less than 0.8) and nuclei that could be classified in another component (log-likelihood probability greater than 0.2) (see Methods). See Supplemental Data Set 1 for exact values. **(E)** The mean of the normalized Hoechst intensity (purple squares, left axis) and projected nuclear area (orange circles, right axis) with the fold increase to the next ploidy class using the 2D Gaussian mixture as shown in (C). **(F)** Segmented nuclear image colored with the ploidy distribution predicted from iSPy shown in (C) using the images from (A).

We observed a wide range of values for the normalized Hoechst intensity and projected nuclear area, regardless of the severity of injury (Fig. 3B). Overall, the distribution of nuclei from the uninjured pylorus had a lower Hoechst intensity and projected area than those from either the mild or severe injuries. We also observed a clear distinction between the distribution of nuclei from the mildly and severely injured pylori. From previous work, we expected to identify nuclei with a ploidy between 2C and 16C^46^. Therefore, we filtered out any nuclei that had a normalized Hoechst intensity of less than 2^-0.5^, because these represented nuclei with a ploidy of less than 1C (Fig. 3B, dashed white line; Methods).

Similar to the analysis of the Arabidopsis data, we compared the performance of three different Gaussian mixtures: a 1D Gaussian mixture using the normalized Hoechst intensity (Fig. S7A–E), a 1D Gaussian mixture using the projected nuclear area (Fig. S7F–J), and a 2D Gaussian mixture using both of these features (Figs. 3C and S8). For the 1D Gaussian mixtures, five components were optimal when using AIC, and four components were optimal when using BIC (Figs. S7A,F). When the curves were assessed by eye, five components better fit the data than four components, which led to one much larger Gaussian curve that encompassed two ploidy classes (Figs. S7D–E, I–J, arrows). Therefore, we focused our analysis on the 1D Gaussian mixtures with five components, corresponding to the following ploidy classes: ≤2C/Unclassified, 2C, 4C, 8C, and ≥16C, because some nuclei had such a high concentration of normalized Hoechst intensity that they may be higher than 16C. Given that the 1D Gaussian mixture identified five components as optimal, we searched for the best fit for the 2D Gaussian mixture using one to five components. Both AIC and BIC predicted that five components provided the best fit (Figs. 3C, S8B–C; Methods).

As a result of the classifications by iSPy, we found a marked difference among the ploidy distributions of the three different pylori (Figs. 3D, S7C,H, and S8D). Our analysis of pyloric ploidy after mild and severe injury recapitulated previous results, with the majority of uninjured pyloric nuclei being 2C, the majority of mildly injured pyloric nuclei being 4C, and the majority of severely injured pyloric nuclei being ≥16C^43^. However, iSPy showed there was a mixture of ploidy in each tissue because uninjured tissues had nuclei between 2C and 8C but both mild and severely injured tissues had nuclei between 2C and ≥16C. We also found that the mean normalized Hoechst intensity increased 1.71–1.91-fold as ploidy doubled, except for the ≥16C group, which increased 3.06-fold from the 8C group, providing more evidence that there may be nuclei that are ≥16C in this group (Fig. 3E, S8E). After this classification, iSPy output maximal projections showing the spatial arrangement of nuclear ploidy in the uninjured, mildly injured, and severely injured pylori (Fig. 3F). These color-coded maps reveal that pyloric nuclei with increased ploidy exist throughout the anterior-posterior axis of the tissue for both the mild and severe injury cases. Therefore, we found that the iSPy method recapitulates the injury severity-dependent endopolyploidy in the Drosophila pylorus while providing insights into the distributions of ploidy throughout the regenerated tissue.

### iSPy verifies tissue-specific polyploidy in human cardiomyocytes

Lastly, we assessed the ability of iSPy to analyze ploidy in tissue sections from larger organs, using the human heart as an example. The muscle cells of the heart, known as cardiomyocytes, become endopolyploid throughout the development of many organisms^49,50^. Developmental cardiomyocyte endopolyploidy is conserved across many living organisms, making it an appropriate polyploidy system to study^7,51,52^. Following developmental polyploidization, ploidy in the adult human myocardium remains stable^53^. Recently, we showed that in adult human cardiomyocytes, the degree to which cardiomyocytes become polyploid during development is chamber-specific and correlates with higher levels of insulin signaling^7^. This chamber specificity leads to higher cardiomyocyte ploidy in the adult left ventricle (LV) relative to the left atrium (LA).

In this study, we re-analyzed confocal microscopy images of frozen heart tissue sections taken from anonymized donors (men aged between 41 and 44 from the Duke Human Heart Repository (DHHR), see Methods). The tissues were labeled with Hoechst to analyze DNA content, wheat germ agglutinin (WGA) to mark cell walls, and phalloidin to mark actin, and imaged via fluorescent confocal microscopy (Fig. 4A, Methods). We performed a sum projection and segmented the resulting two-dimensional image in ilastik. Using the phalloidin labeling of cardiomyocyte muscle striations as a guide, we manually selected the cardiomyocytes from each image and extracted the total intensity of the Hoechst staining and the projected nuclear area (Fig. 4A, yellow nuclei, see Methods).

**Figure 4:**
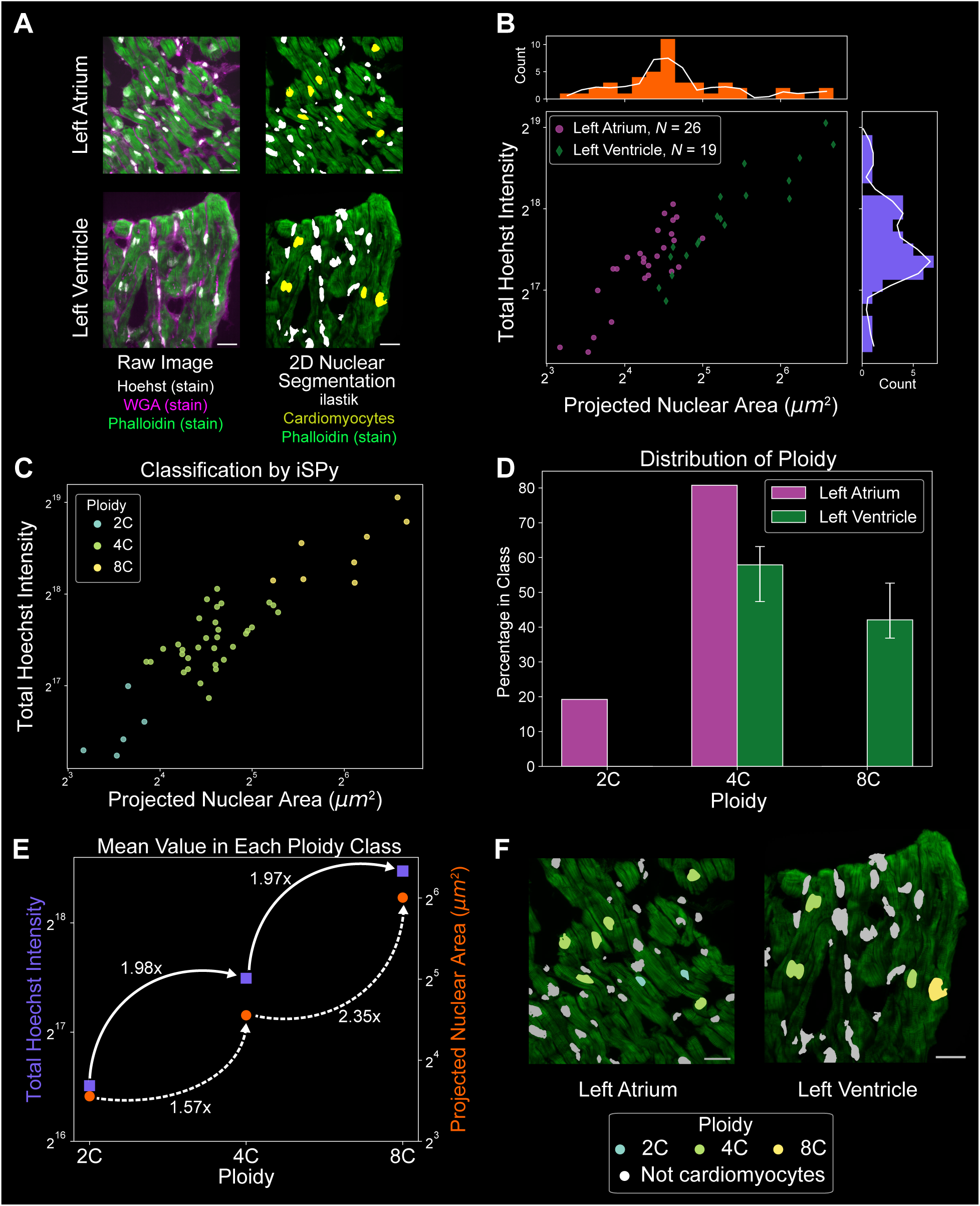
iSPy predicts three distinct ploidy classes for human cardiomyocytes. **(A)** Left: representative sum-projections of confocal images of human heart tissue stained with Hoechst (white), wheat germ agglutinin (WGA, magenta), and phalloidin (green) from the left atrium (top) and left ventricle (bottom). Right: segmented nuclei using ilastik. Yellow nuclei signify hand-selected cardiomyocytes using phalloidin as a guide (see Methods). Note this segmentation was performed with the sum projection in two dimensions. Scale bars: 25 µm. **(B)** Scatter plot of the projected nuclear area and total Hoechst intensity of the segmented cardiomyocytes (left atrium, purple circles; left ventricle, green diamonds) including corresponding histograms of the normalized Hoechst intensity (purple) and projected nuclear area (orange) with a smoothed Savitzky–Golay filter (white line, only for illustrative purposes). **(C)** iSPy ploidy prediction using the 2D Gaussian mixture with 3 components and spherical covariance matrix (see Fig. S9A for additional information). **(D)** The proportion of cardiomyocytes in each ploidy class predicted from the 2D Gaussian mixture in (C) by heart chamber (left atrium, purple; left ventricle, green). Uncertainty bars represent nuclei that may be classified incorrectly (log-likelihood probability less than 0.8) and nuclei that could be classified in another component (log-likelihood probability greater than 0.2) (see Methods). See Supplemental Data Set 1 for exact values. **(E)** The mean of the Hoechst intensity (purple squares, left axis) and projected nuclear area (orange circles, right axis) with the fold increase to the next ploidy class using the 2D Gaussian mixture from (C). **(F)** Segmented nuclear image colored with the ploidy distribution predicted from the 2D Gaussian mixture (C) using the images from (A). Scale bars = 25 µm.

As we found previously using individual hand-drawn nuclear segmentation, we found that LV cardiomyocytes had a larger projected nuclear area and a higher total Hoechst intensity than the LA cardiomyocytes (Fig. 4B)^7^. We observed clusters with varying levels of Hoechst intensity and nuclear area, similar to previous findings, which showed that cardiomyocytes are between 2C and 8C^49^. Thus, we expected to find three classes of ploidy: 2C, 4C, and 8C and this is indeed what we observed using iSPy.

Due to the relatively small sample size (*N* = 26 for LA, *N* = 19 for LV), we only performed a 2D Gaussian Mixture with both total Hoechst intensity and the projected nuclear area. We searched for the best fit by varying the number of components between one and three and found that three components provided the best fit for both information criteria and for either spherical or diagonal covariance matrices (Figs. 4C, S9A–C). As expected from our previous work, the ploidy proportion of LV and LA cardiomyocytes differed greatly, with LA cardiomyocytes having a ploidy of 2C and 4C and LV cardiomyocytes having a ploidy of either 4C or 8C (Fig. 4D, S9D). Moreover, our procedure accurately captured the 2-fold increase in total Hoechst intensity as ploidy increased whereas we observed a 1.6–2.5-fold or 2–2.8-fold increase in the projected nuclear area as ploidy increased, depending on the covariance matrices used (Fig. 4E, S9E). Following this classification, iSPy generated a maximal projection image showing the spatial arrangement of nuclear ploidy of the classified cardiomyocyte ploidy from our predictions (Fig. 4F). Thus, iSPy verified a chamber-specific ploidy dependence in human cardiomyocytes using the Hoechst intensity and projected nuclear area.

## Discussion

Inferring the ploidy of nuclei within a tissue without destroying the integrity of the sample is critically important for the study of the development and growth of an organism and also for the analysis of emerging spatial patterns related to ploidy. In this paper, we introduce iSPy, a high-throughput method of measuring the ploidy of nuclei while simultaneously maintaining the spatial integrity of the tissue samples. This method uses simple experimental and segmentation tools along with an unsupervised Gaussian mixture model that uses only the size of the nuclei and the intensity of the fluorescence/staining signals to accurately identify nuclear ploidy and output a maximal projection image of the segmentation with its ploidy prediction. Table 1 summarizes the techniques used with iSPy experimentally and computationally for each model organism discussed in this work. Based on our analysis presented here, our pipeline could be applied to all tissue types in any organism, regardless of tissue properties or the need to perform tissue sectioning. The inclusion of other nuclear or cell membrane markers would allow detailed analyses regarding the effects of gene expression and cell size on ploidy. iSPy can also be effective in live-imaging and time-lapse experiments, analyzing ploidy development both in time and space across a tissue.

**Table 1:** Methodology and results summary for iSPy on Arabidopsis cotyledons, Drosophila pylorus, and human heart tissue. See the Methods section and Results sections for more specific details.

| Tissue | Experiments | ilastik Segmentation | iSPy Analysis |
| --- | --- | --- | --- |
| *Arabidopsis thaliana* cotyledon | – Nuclear marker: *p35S::H2B-RFP1*<br>– Live or fixed tissue | – Three-dimensional volume segmentation<br>– Hand-select stomatal guard cells and remove incomplete nuclei | – Remove sub-epidermal cells using a threshold for "Variance of Intensity"<br>– 2D iSPy using nuclear volume and total H2B-RFP1 intensity: 6 components, spherical or diagonal covariance<br>– Remove nuclei in the component with the lowest mean<br>– 2–2.4-fold increase in total intensity, ~1.8-fold increase in volume |
| *Drosophila melanogaster* pylorus | – Protocol from Clay et al. (2023)<br>– Nuclear marker: Hoechst staining | – Two-dimensional segmentation using sum projection<br>– Hand-select and remove incomplete nuclei | – Normalize pyloric nuclei by 1C sperm nuclei<br>– 1D iSPy using normalized Hoechst intensity, 5 components, full covariance using AIC criterion<br>– 2D iSPy using normalized Hoechst intensity and projected area: 5 components, spherical or diagonal covariance<br>– 1.7–1.9-fold increase in normalized intensity, ~1.5 fold increase in projected nuclear area |
| Human heart | – Protocol from Chakraborty et al. (2023)<br>– Nuclear marker: Hoechst staining<br>– Optional staining: WGA and phalloidin to help identify cardiomyocytes | – Two-dimensional segmentation using sum projection<br>– Hand-select cardiomyocytes (using phalloidin to help identify them) | – 2D iSPy using total Hoechst intensity and projected nuclear area: 3 components, spherical or diagonal covariance<br>– 2-fold increase in total Hoechst intensity, 1.5–2.5-fold increase in projected nuclear area |

Some improvements can be made to this technique in the future. Other histone markers and promoters could be examined to see whether better candidates exist for ploidy quantification. Although unsupervised Gaussian mixture models can easily identify well-separated clusters, other supervised and unsupervised clustering algorithms can be tested, especially if there are systems where assuming Gaussian distributions is not possible (e.g., k-means and hierarchical clustering algorithms)^54^. Furthermore, these clusters might be easier to identify when more nuclear geometrical features are taken into account, and the implementation of spatial statistics such as radial distribution functions or tools from topological data analysis might allow patterns and underlying ploidy architecture to be identified^55^.

In this study, we employed ilastik as a segmentation software, but other segmentation tools are available and can be used with our software, such as Fiji, CellPose, PlantSeg, CelloType, and MorphoGraphX^56–59^. Some of these tools can be high-throughput, but they may have a steeper learning curve than ilastik, particularly for those users who are not familiar with segmentation software. However, precision and speed should be balanced with user-friendliness when conducting segmentations, particularly for large data sets. Therefore, when considering whether to use ilastik, one should consider the number of tissue samples involved. For example, in this study in which we are limited by the number of tissue sections from human heart samples, the amount of effort did not save more time than our previous approach where we individually identified each nucleus by eye^7^. However, as one scales up to larger numbers of samples and images, the high-throughput capabilities of ilastik become superior to those of the individual nucleus identification approaches.

In conclusion, this *in silico* methodology opens a new avenue to assess – in a high-throughput manner – how ploidy affects nuclear size, cells, and tissues, and how endopolyploidy is spatially patterned across organisms.

## Methods

### Plant growth conditions

For flow cytometry, seeds were sown directly onto Lambert LM-111 All Purpose Mix soil and stratified at 4°C for 2 days in darkness. Plants were grown under continuous fluorescent light (∼80 μmol m−2 s−1) at 22°C and 60–75% relative humidity. The Columbia-0 (Col-0) accession line carrying *pML1::mCitrine-RCI2A* and *pML1::H2B-mTFP* was used^37,40^. Cotyledons were harvested 14 days after stratification. For confocal imaging, seeds were stratified in water for 3-4 days at 4°C before being sown onto soil. Plants were grown in a Percival AR-95L3 chamber at 22°C and 60% relative humidity at 15% light (4592 Lux) in continuous light. The Col-0 accession line carrying *p35S::H2B-RFP1* and *pUBQ10::MYR-CFP* was used. This line was generated by Weibeing Yang, by transforming the membrane marker *pUBQ10::MYR-CFP* into Col-0 wild-type plants and afterwards crossing them with Col-0 plants expressing *p35S::H2B-RFP1*^60^. Cotyledons were harvested 14 days after stratification.

### Flow cytometry for Arabidopsis thaliana

Tissue was harvested from 6–8 cotyledons per sample at 14 days post-stratification. Flow cytometry was performed as described previously^37,61,62^. For each sample, harvested tissue was thoroughly chopped with a sterile razor blade in a Petri dish on ice containing 470 μL Aru buffer (10mM MgSO4 hydrate, 50mM KCl, 5mM HEPES, 10mM DTT (dithiothreitol), 2.5% v/v Triton X-100). The solution was filtered through a 40-μm Fisherbrand cell strainer and 350 μL was transferred to a 5 mL round bottom tube. Suspended nuclei were then treated with RNAse (0.1 mg / 100 μL sample), and stained with PI (0.001 mg / 100 μL sample). Samples were run on a BD Accuri C6 flow cytometer. Events were gated to separate epidermal (TFP-positive) nuclei from non-epidermal (TFP-negative) nuclei using the FL1 (533/30) channel. Relative nuclear DNA content was determined by PI fluorescence of epidermal and non-epidermal cells using the FL2 (585/40) channel.

### Data acquisition for *Arabidopsis thaliana*

For confocal imaging, a Leica Stellaris 8 confocal microscope was used, with a NA 0.95 and 25× water dipping objective and a z-step of 0.4 μm. Cotyledons were placed onto microscopy slides with a cover slip. To detect *p35S::H2B-RFP1*, a laser with 590 nm emission wavelength and 2.0 intensity was used, and fluorescence was captured at 605–644 nm with a 45.5% gain. For *pML1::H2B-mTFP*, a laser with 462 nm emission wavelength and 2.0 intensity was used, and fluorescence was captured at 467–520 nm with a 20% gain. For *pML1::mCitrine-RCI2A*, the laser emission wavelength was 515 nm, laser intensity was set to 2.0 intensity, and fluorescence was captured at 523–556 nm with a 47.8% gain. Following acquisition, image files were stitched together inside LAS X software and then exported. The adaxial and abaxial sides of two cotyledons were imaged.

### *Drosophila melanogaster* experimental conditions and data acquisition

All female adult flies were aged for 4–7 days after eclosion before injury experiments. Flies were kept at 18**°**C except during injury. Injury through a tissue-specific, genetic ablation system was induced in a temperature-dependent manner; flies were transferred from an 18°C to a 29°C incubator to induce injury. The class of severity of injury was determined by the duration that the flies were at 29**°**C; mildly injured flies were kept at 29°C for 24 h, while severely injured flies were kept at 29**°**C for 48 h. Flies were allowed to recover for at least 3–4 days before dissection. As described previously, *byn-Gal4, tub-Gal80^ts^, UAS-hid*/*TM3 Sb* was used for injured and uninjured flies^43^. For each injury condition, 3–4 pylori and 2 Drosophila testes (the sperm of which are 1C haploid control) were placed on the same slide^26^. Details of the tissue sample preparation and Hoechst staining have been previously described^26^. Samples were imaged using a Nikon Ti2 Eclipse with a Nikon A1 camera and an NA 1.42 and 60× oil dipping objective (step size: 0.4 µm). To detect Hoechst, the laser emission wavelength was 405 nm with 1.78 intensity, and fluorescence was captured at 420–480nm with 42% gain.

### Cardiomyocyte experimental conditions and data acquisition

Human heart tissue samples from explanted hearts were obtained from the Duke Human Heart Repository (DHHR) with approval from the Duke University Health System (DUHS) Institutional Review Board (IRB) (Pro00005621). Details of the tissue sample preparation, immunostaining, and imaging have been previously described^7^. Briefly, flash-frozen human left ventricular (LV) and left atrial (LA) tissue samples were sectioned at a thickness of 10 µm and immunostained with wheat germ agglutinin (WGA) (1:250, W21404; Invitrogen), Alexa Fluor® 488 Phalloidin (1:250, 8878; Cell Signaling), and Hoechst. Samples were imaged using an Andor Dragonfly 505 system with Borealis illumination on a spinning-disk confocal microscope (z-step size: 0.5 µm) and an Andor Zyla PLUS 4.2 Megapixel sCMOS camera, coupled with a 63×/1.47 TIRF HC PL APO CORR (Leica 11506319) oil objective (working distance: 0.10 mm).

### Nuclear segmentation and data processing for *Arabidopsis thaliana*

After image acquisition, images (as .lif files) were imported into Fiji (ImageJ) and exported as HDF5 files (.h5) using the ilastik plugin. Additionally, three crops of each confocal image were exported as an HDF5 to train ilastik with. In ilastik, the “Pixel + Object Classification Workflow” was used to import the three cropped images. After conducting the pixel training between nuclei and background on all three crops, hysteresis thresholding was performed for the nuclear segmentation with smoothing parameters (1.5, 1.5, 1.5), a core threshold of 0.75, and a final threshold of 0.45, while marking the “Don’t merge objects” box. All available nuclear features were calculated and no nuclei were classified during the “Object Classification” step. Object Predictions were exported using a 16-bit data type as an HDF5 file as well as the CSV file for the Feature Table. Batch Processing was performed on the large file corresponding to the three individual crops, which automatically output the Object Predictions HDF5 file and the CSV file with nuclear features. Then the “Object Classification [Inputs: Raw Data, Segmentation]” Workflow (creating a whole new ilastik file) was used and both the raw data file and the segmentation file of the large image were imported. Stomatal guard cell nuclei and nuclei that were incorrectly segmented, merged with another nucleus, or not completed (nuclei on the borders) were hand-selected. The Object Predictions and Object Identities were exported along with the CSV file, which was updated with the correct labels in the “User Labels” column. The “Predicted Class” column can also be used if ilastik’s neural network function is used to classify certain nuclear types.

After nuclei segmentation, sub-epidermal nuclei were filtered out using the “Variance of intensity” quantity. This calculates the inhomogeneity of the fluorescence signal across each nucleus. Mathematically, the variance of intensity σ_*i*_ for each nucleus *i* is calculated as:

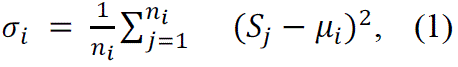

where *n*_*i*_ is the number of voxels that correspond to the segmented nuclear volume of the nucleus *i*, *S_j_* is the intensity of the fluorescent marker at pixel *j* for nucleus *i*, and μ_*i*_ is the mean intensity of the fluorescent marker of nucleus *i*. Plotting the variance of pixel intensity in each nucleus for all nuclei gave two large peaks (Fig. S3B). A threshold of 2^8.3^, which lies right in the trough of the two peaks, was used (dashed white line, Fig. S3B). Sub-epidermal nuclei were defined to have a variance of intensity lower than this threshold. From our observations, these nuclei are sub-epidermal (Fig. S3C). For the flow cytometry data, nuclei were filtered out that had a total *pML::H2B-TFP* intensity lower than 2^8^ because these nuclei are most likely stomatal guard cell nuclei (Fig. S2F).

### Nuclear segmentation and data processing for *Drosophila melanogaster*

Following image acquisition, images were imported into Fiji/ImageJ and converted to a sum projection. This was then exported as a TIF file. The “2D Pixel Classification” workflow was used to create a pixel probability map for each image. After exporting the probability map, the “Object Classification [Inputs: Raw Data, Pixel Prediction Map]” workflow was used to conduct hysteresis thresholding for the nuclear segmentation using the smoothing parameters (1.0, 1.0), a core threshold of 0.6–0.8, and a final threshold of 0.5–0.7, depending on the image. All available nuclear features were calculated except “Skewness of intensity”, “Skewness of Intensity in neighborhood”, “Kurtosis of Intensity”, and “Kurtosis of Intensity in neighborhood”. Then, for the uninjured and mild injury pylori, pyloric nuclei were manually selected that were either incomplete (e.g., partially out of frame) or belonged to a neighboring tissue and removed from quantification. For the severely injured pylori and spermatids, all nuclei/spermatids that were complete and belonged to the tissue were selected by hand. The Object Predictions and Object Identities were exported along with the CSV file, which was updated with the correct labels in the “User Labels” column.

The total Hoechst intensity (column name “Total Intensity” or “Total Intensity_0”, depending on the version of ilastik) was normalized by the spermatid intensities as in previous work^26^. For each experiment, spermatids were imaged on the same slide as the fly tissue. Only those experiments in which we obtained at least 10 spermatids were used, where the median spermatid intensity was 2,000–6,000, and 90% of the spermatid nuclear intensity – when divided by the median spermatid intensity – was 0.5–1.5 (see Fig. S6B). For experiments that met these conditions, the intensity of each pyloric nucleus was divided by the median spermatid intensity to create a “Normalized Hoechst Intensity” quantity, whereby a value of 2^0^ = 1 corresponded to 1C, 2^1^ corresponded to 2C, 2^2^ to 4C, and so on. Any experiments that did not satisfy the spermatid criteria were not considered.

### Nuclear segmentation and data processing for human cardiomyocytes

Microscopy images were exported as Imaris File format files (.ims) and the nuclei were visualized in three dimensions using the ImarisCell module where cardiomyocyte nuclei that were complete and not partially severed were identified (Imaris Version 8.2). Cardiomyocyte nuclei were selected using both phalloidin and WGA labeling. The .ims files were imported into Fiji and were exported as TIF files for ilastik analysis. Not all organ donor samples were analyzed. The “2D Pixel Classification” workflow was used to create a pixel probability map for each image. Then, the “Object classification [Inputs: Raw Data, Pixel Prediction Map]” workflow was used to conduct hysteresis thresholding for the nuclear segmentation with the smoothing parameters (1.0, 1.0), a core threshold of 0.8, and a final threshold of 0.5. All available nuclear features were calculated. When using the Object Classification step, cardiomyocytes with complete nuclei – verified by the 3D rendering in ImarisCell – were labeled as “Cardiomyocyte nucleus” and were used for further analysis. The Object Predictions and Object Identities were exported, along with the CSV file, which was updated with the correct labels in the “User Labels” column.

### Gaussian mixture models

All ploidy predictions were performed using unsupervised Gaussian mixture models in the Scikit-learn Python package^34^, which implements the expectation-maximization algorithm^63^. In this work, either one-or two-dimensional Gaussians (1D or 2D) were used, depending on whether one (e.g., total signal intensity) or two features (e.g., total signal intensity and nuclear volume) were considered. Before performing the Gaussian mixture model, the data were logarithmically scaled by 2, which implied that each Gaussian would have a probability density function similar to a log-normal distribution with base 2. Although this is an unsupervised learning algorithm, the optimal fit was searched for by performing Gaussian mixture models while varying the number of components (i.e., the number of Gaussians added together to make the full Gaussian mixture) and the types of covariance matrices that were available in the Scikit-learn package: spherical (each component has a single variance), diagonal (each component has its own diagonal covariance matrix), full (each component has its own general covariance matrix), and tied (all components share the same general covariance matrix)^34^. We note that the spherical covariance matrix with 5 components was used to compute the 1D Gaussian mixture for the *pML::H2B-TFP* intensity.

For the 1D Gaussian mixtures, a tolerance of 10^-5^, a maximum number of iterations of 10,000, 20 initializations, and the k-means++ initialization method were used. For the 2D Gaussian mixtures, a tolerance of 10^-7^, a maximum number of iterations of 5,000, 500 initializations, and the k-means++ initialization method were used. For the 1D Gaussian mixture for the PI staining intensity, the spherical covariance matrix with 5 components with 10^-7^, maximum iterations of 5,000, 500 initializations, and k-means++ initialization method was used (Fig. S2G). Such parameter values were set to avoid underfitting the data.

After identifying the optimal fit for each combination of components and covariance matrices, both the Akaike and Bayesian Information Criteria (AIC and BIC, respectively) were used to find the best fit given a certain covariance matrix^34,35,64^. We use the definition used in the Scikit-learn package, i.e.,

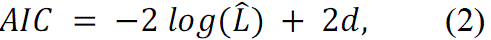

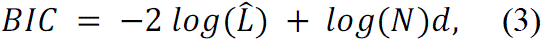

where 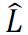 *L* is the maximum likelihood of the model, *d* is the number of parameters, and N is the number of samples (in this case, nuclei)^34^. This model was subsequently used after obtaining the best fit from the information criteria. The predict function was then used, which assigns which component each nucleus is likely to belong to by using the log-likelihood probabilities.

### Statistics

To create the uncertainty bars for all histograms, the following procedure was utilized where *N* is the number of samples and *K* is the number of components computed from the Gaussian mixture model:

1. The log-likelihood probabilities 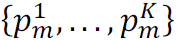 for each nucleus *m* = {1, …, N} to belong to each component *k* = {1, …, *K*} were computed using the predict_proba function. Note that 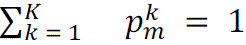 for all *m*.
2. The maximum log-likelihood probability for a nucleus *m* was defined as 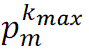. This implies that the nucleus *m* was classified as part of the component *k_max_*,. The proportions of nuclei that were classified into component *k*, i.e. all *m* such that *k_max_*,= *k*, were defined as the set {*V*^1^, …, *V^K^*}.
3. If 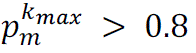, the classification was confident enough that *m* was classified into the component *k_max_*.
4. The number of nuclei classified into each component where 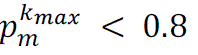 was calculated and defined 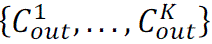 as the proportion of nuclei that could be classified to another component.
5. Lastly, all nuclei in which the second-largest log-likelihood probability was greater than 0.2 were found. The number of nuclei that satisfied this condition for each component was calculated and 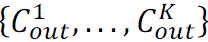 was defined as the proportion of nuclei that could be classified into one of these components.
6. Therefore, with the above considerations, the uncertainty bars were defined as those which spanned the interval 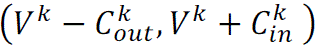 H for each component *k*.

## Supporting information

Supplemental Data Set 1

## Acknowledgments

We thank Weibing Yang for generating the line carrying the reporters *pUBQ10::MYR-CFP* × *p35S::H2B-mRFP1* and kindly sharing it with us. Note the membrane marker he created is not shown in the current publication. We thank John Chandler, Franziska Turck, Jake Klemm, Chun-Biu Li, and André Marques for providing critical feedback on the manuscript. Funding for this work is acknowledged from the Max Planck Society for N.J.R. and P.F.J.; the Deutsche Forschungsgemeinschaft (DFG, German Research Foundation) under Germany’s Excellence Strategy – EXC-2048/1 – project ID 390686111 for P.F.J; the Polyploidy Integration and Innovation Institute (PI3) US NSF DBI-2320251 for D.F., P.F.J, and A.H.K.R; and NIH grant R01GM118447 for D.T.F.

## Author Contributions

Conceptualization: N.J.R., P.B.B., L.S.O., P.F.J., D.T.F.; Data curation: N.J.R., P.B.B., L.S.O., A.C.; Formal analysis: N.J.R., P.B.B., L.S.O., A.C.; Funding acquisition: P.F.J., D.T.F., A.C., A.H.K.R.; Investigation: N.J.R., P.B.B., L.S.O., A.C.; Methodology: N.J.R., P.B.B., L.S.O., A.C.; Project administration: N.J.R., P.F.J., D.T.F.; Resources: P.F.J.; Software: N.J.R.; Supervision: P.F.J. D.T.F., A.H.K.R.; Validation: N.J.R., P.B.B., L.S.O., A.C.; Visualization: N.J.R.; Writing – original draft: N.J.R., D.T.F., P.B.B., A.C., L.S.O.; Writing – review & editing: N.J.R., P.F.J., D.T.F., P.B.B., A.H.K.R., L.S.O.

## Data availability

All data are available in the main text, in the supplementary materials, and are publicly available in our OSF data repository https://osf.io/um7r3/. The GitLab repository for this work is here: https://gitlab.gwdg.de/devplantpatterning/Publications/ispy-inferring-spatial-ploidy.

