## Supplemental Figures for "Spatial ploidy inference using quantitative imaging"

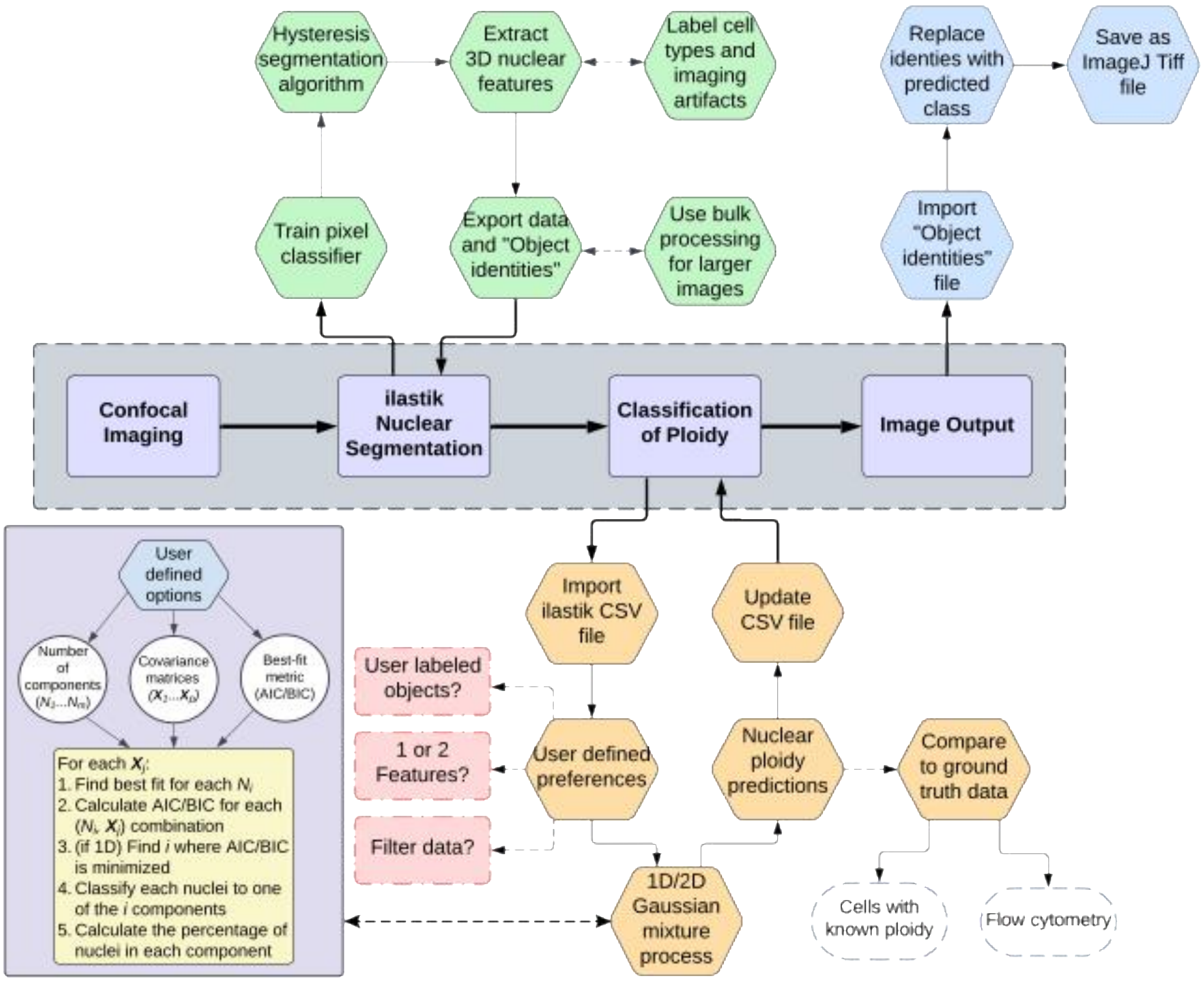


**<u>Figure S1: Extended version of Fig. 1B.</u>** The full iSPy methodology that includes the Gaussian mixture implementation. See Results and Methods for further details.

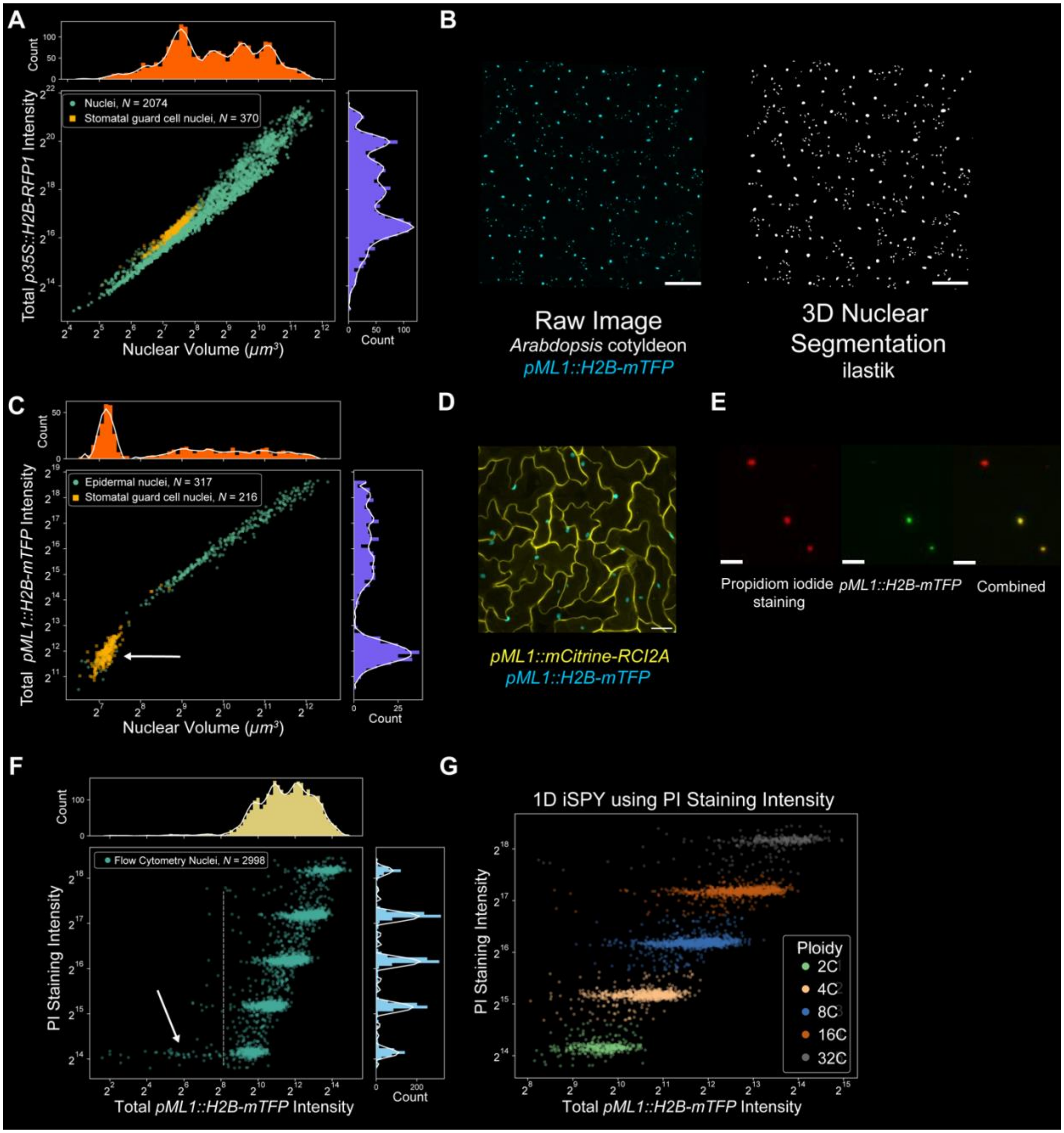


**<u>Figure S2: *pML1::H2B-mTFP* can identify epidermal cells and stomatal guard cells.</u>** **(A)** Scatter plot of the nuclear volume and total *p35S::H2B-RFP1* intensity of the ilastik-segmented nuclei (green) and stomatal guard cell nuclei (yellow) including corresponding histograms of the total *p35S::H2B-RFP1* intensity (purple) and nuclear volume (orange) with a smoothed Savitzky–Golay filter (white line, only for illustrative purposes). **(B)** Left: representative confocal image of *pML1::H2B-mTFP* in a 14-day-old cotyledon. Right: nuclear segmentation of the left image performed in ilastik. Note that segmentation was performed in three dimensions. Scale bars = 100 µm. **(C)** Scatter plot of the nuclear volume and total *p35S::H2B-RFP1* intensity of the segmented nuclei (epidermal nuclei, green circles; stomatal guard cells, yellow squares) including representative histograms of the total *p35S::H2B-RFP1* intensity (purple) and nuclear volume (orange) with a smoothed Savitzky–Golay filter (white line, only for illustrative purposes).

Note the cluster of stomatal guard cells (arrow) and the lack of clusters in the epidermal cells, contrary to what was observed with *p35S::H2B-RFP1* (see Fig. 2B, Methods). **(D–G)** Data from flow cytometry (see Methods for experimental information). **(D)** Representative image of a 14-day cotyledon used for flow cytometry tagged with epidermal-specific cell membrane fluorescent marker *pML1::mCirtrine-RCI2A* (yellow) and the epidermal-specific nuclear marker *pML1::H2B-mTFP* (blue). Scale bar = 25 µm. **(E)** A confocal image of nuclei from a *pML1::mCirtrine-RCI2A* × *pML1::H2B-mTFP* cotyledon stained with propidium iodide (PI) (left). Note that not all cells express *pML1::H2B-mTFP* (middle, right), which allows the identification of epidermal cells. Scale bars: 20 µm. **(F)** Scatter plot of the total *pML1::H2B-mTFP* intensity and total PI staining intensity including representative histograms of the total *pML1::H2B-mTFP* intensity (yellow) and total PI staining intensity (blue) with a smoothed Savitzky–Golay filter (white line, only for illustrative purposes). Only cells that express *pML1::H2B-mTFP* expression are shown (i.e., epidermal cells). Note the cluster of stomatal guard cells (arrow) similar to the confocal imaging data in (C). We use a threshold of $2^8$ to remove all stomatal guard cells from the calculations (see Fig. 2). **(G)** Classification of ploidy from the flow cytometry data using a 1D Gaussian mixture with five components and full covariance matrices on the total intensity of the PI staining. Percentages of nuclei in each ploidy class can be found in Fig. 2D (green bar).

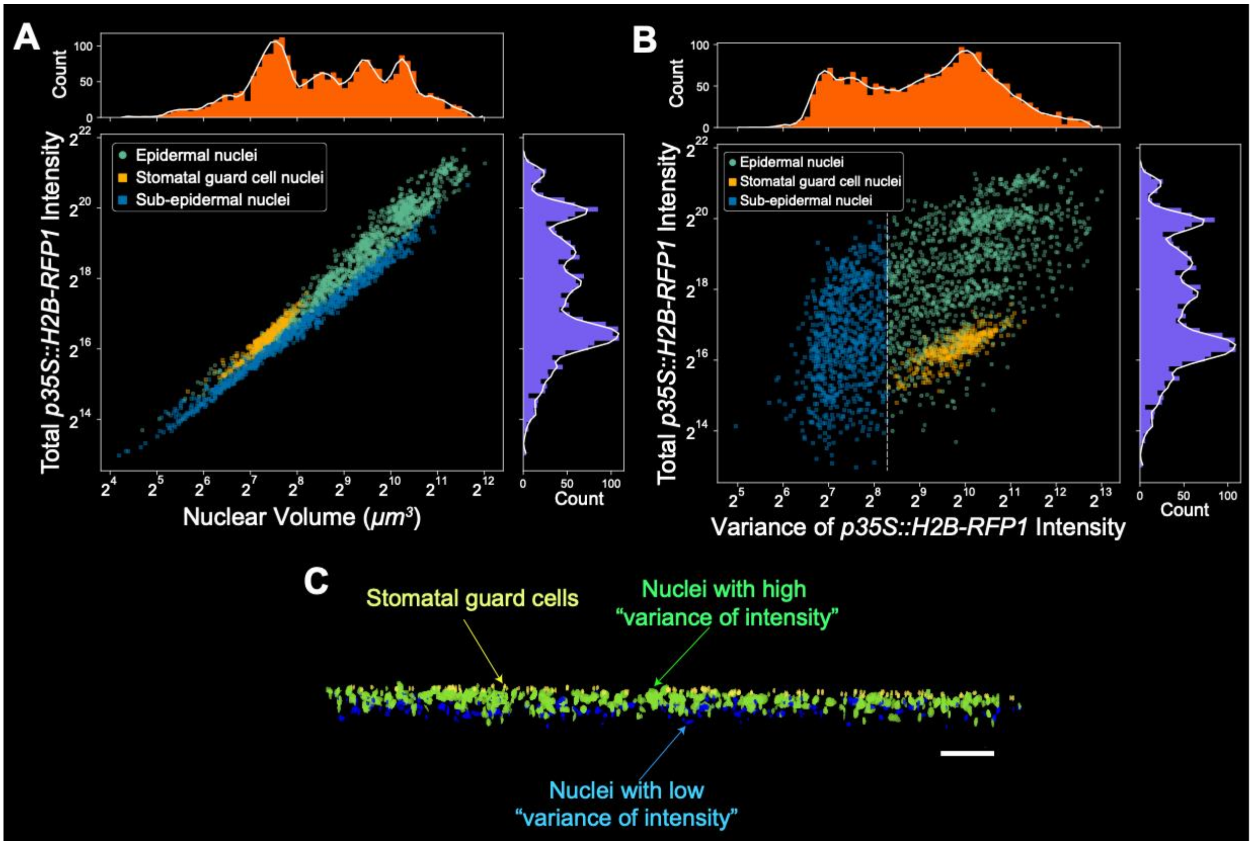


**Figure S3: Sub-epidermal cells have a low variance of intensity.** **(A)** Scatter plot of the nuclear volume and variance of the *p35S::H2B-RFP1* intensity of the segmented nuclei including representative histograms of the total *p35S::H2B-RFP1* intensity (purple) and variance of the *p35S::H2B-RFP1* intensity (orange) with a smoothed Savitzky–Golay filter (white line, only for illustrative purposes). Note the bi-modal distribution in the histogram of the variance of the *p35S::H2B-RFP1* intensity. A threshold of $2^{8.3}$ (white dashed line) was used and any cells beneath this threshold were classified as sub-epidermal cells (blue squares). **(B)** Scatter plot of the nuclear volume and total *p35S::H2B-RFP1* intensity of the segmented nuclei (epidermal nuclei, green circles; stomatal guard cells, yellow squares; sub-epidermal nuclei, blue squares) including representative histograms of the total *p35S::H2B-RFP1* intensity (purple) and nuclear volume (orange) with a smoothed Savitzky–Golay filter (white line, only for illustrative purposes). **(C)** A representative snapshot of segmented nuclei, with the low-variance of intensity nuclei shaded in blue, and in green and yellow for high-variance of intensity nuclei and stomatal guard cells, respectively. Note that nuclei with a higher variance of intensity are above those with a lower value in the scatter plot. Scale bar = 100 μm.

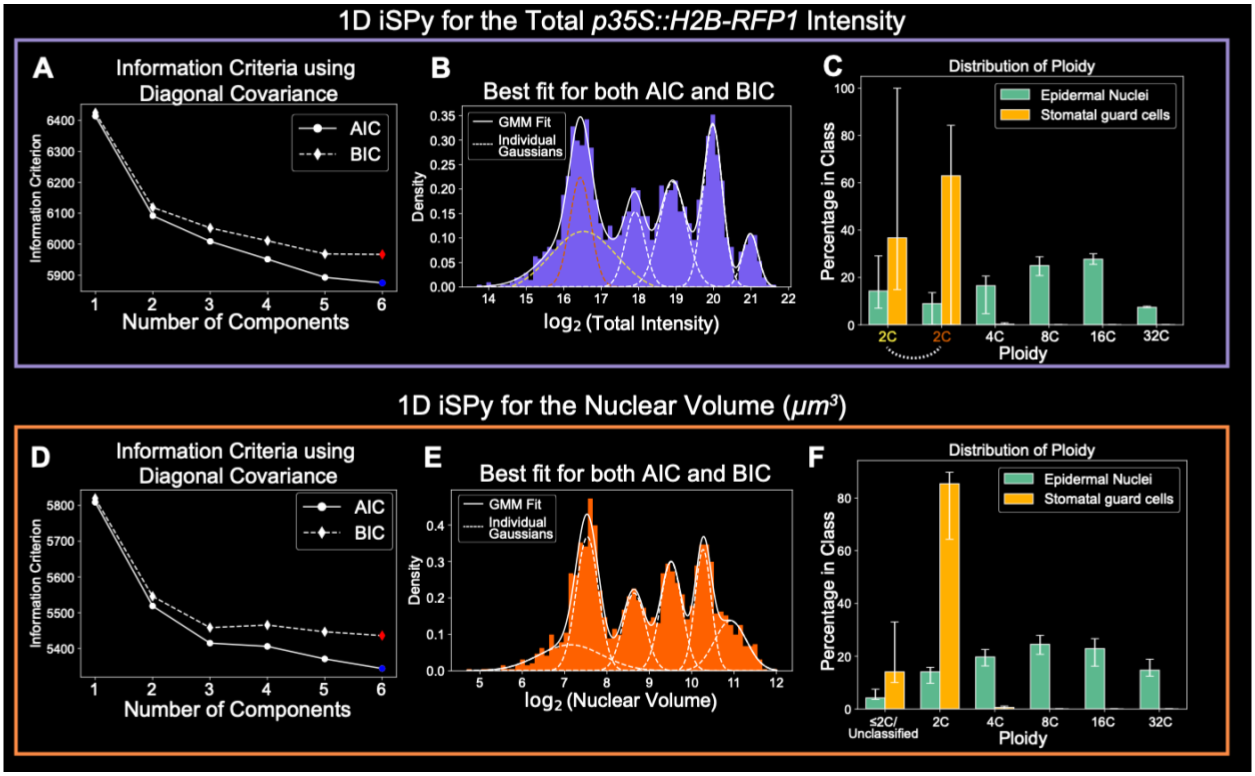


**<u>Figure S4: 1D Gaussian mixtures for Arabidopsis cotyledons predict six components.</u>** **(A–C)** A 1D Gaussian mixture for the total *p35S::H2B-RFP1* intensity (see Methods for details). **(A)** Values of AIC (solid line) and BIC (dashed line) as the number of components is varied while using the diagonal covariance matrices. The blue and red colored points at six components represent the lowest AIC and BIC, respectively, and therefore the best fit. **(B)** The best fit using the 1D Gaussian mixture displaying the histogram (purple), the full Gaussian mixture fit (solid white line), and the six components (dashed lines) that sum up to equal the best fit. The two colored Gaussians (yellow and orange) represent 2C cells, and the subsequent Gaussians represent 4C, 8C, 16C, and 32C cells, respectively. **(C)** The proportion of stomatal guard cells (yellow) and epidermal cells (green) predicted to be in each ploidy class. Note the two 2C classes, which correspond to the colored Gaussians in (B). See Supplemental Data Set 1 for exact values. **(D–F)** A 1D Gaussian mixture for the nuclear volume. **(D)** Values of AIC (solid line) and BIC (dashed line) as the number of components is varied while using the diagonal covariance matrices. The blue and red colored points at 6 components represent the lowest AIC and BIC, respectively, and therefore the best fit. **(E)** The best fit using the 1D Gaussian mixture displaying the histogram (orange), the full Gaussian mixture fit (solid white line), and the six components (dashed lines) that sum up to equal the best fit. The Gaussians represent the unclassified, 2C, 4C, 8C, 16C, and 32C cells, respectively. **(F)** The proportion of stomatal guard cells and epidermal cells predicted to be in each ploidy class. Note the high number of stomatal guard cells in the 2C group compared to the unclassified and 4C groups. See Supplemental Data Set 1 for exact values. Uncertainty bars represent nuclei that may be classified incorrectly (log-likelihood probability less than 0.8) and nuclei that could be classified in another component (log-likelihood probability greater than 0.2) (see Methods).

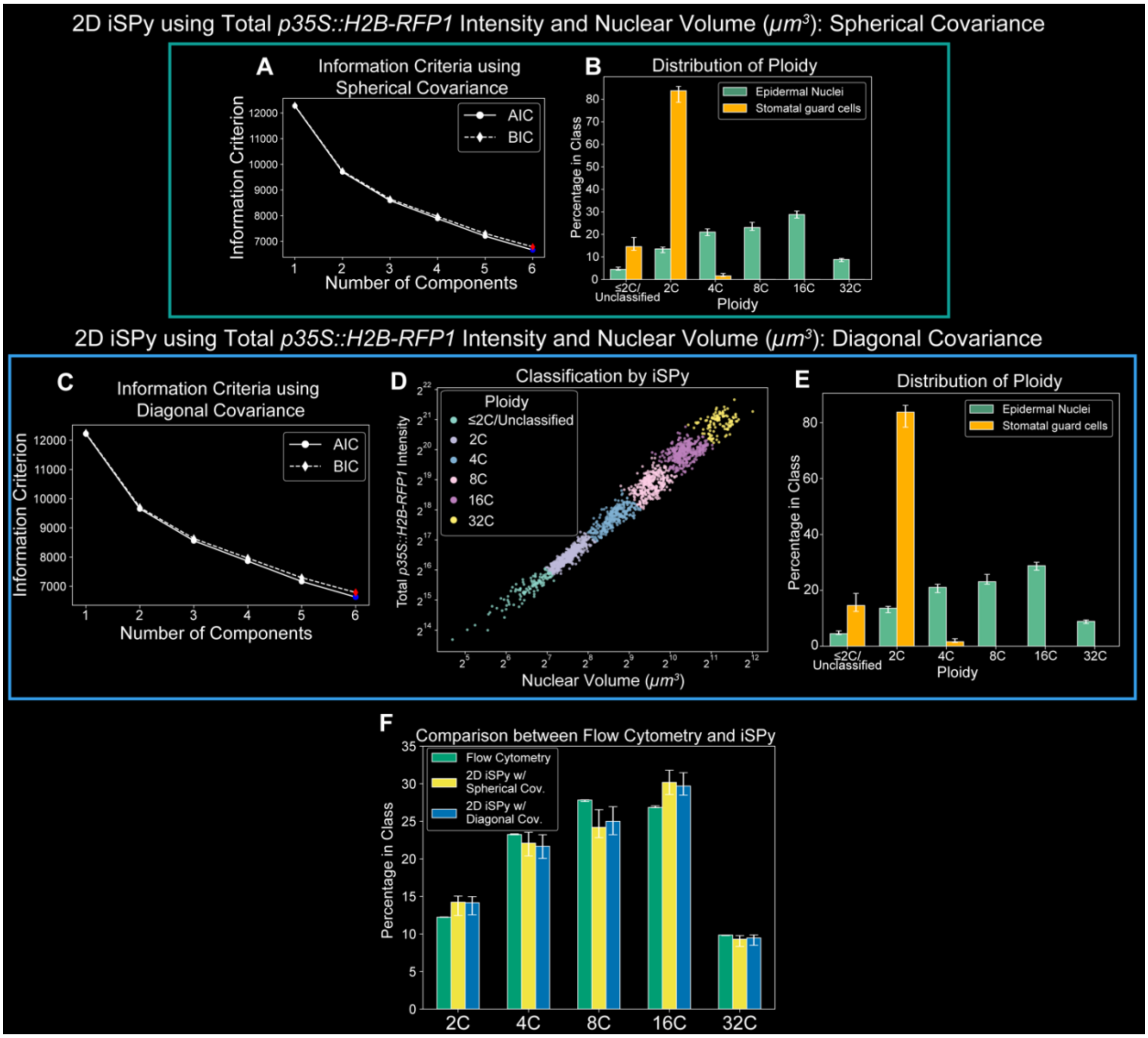


**<u>Figure S5: 2D Gaussian mixtures for Arabidopsis cotyledons predict six components.</u>** **(A–B)** A 2D Gaussian mixture using both total *p35S::H2B-RFP1* intensity and spherical covariance found in Fig. 2C–F. **(A)** Values of AIC (solid line) and BIC (dashed line) as the number of components is varied while using the spherical covariance matrices. The blue and red colored points at six components represent the lowest AIC and BIC, respectively, and therefore the best fit. **(B)** The proportion of stomatal guard cells (yellow) and epidermal cells (green) predicted to be in each ploidy class. **(C–E)** A 2D Gaussian mixture using both total *p35S::H2B-RFP1* intensity and diagonal covariance. See Supplemental Data Set 1 for exact values. **(C)** Values of AIC (solid line) and BIC (dashed line) as the number of components is varied while using the diagonal covariance matrices. The blue and red colored points at 6 components represent the lowest AIC and BIC, respectively, and therefore the best fit. **(D)** iSPy prediction for each epidermal nucleus using the 2D Gaussian mixture with six components and spherical covariance matrix. **(E)** The proportion of stomatal guard cells (yellow) and epidermal cells (green) predicted to be in each ploidy class. **(F)** The comparison of the percentage of epidermal nuclei predicted in each ploidy class between flow cytometry (green, Fig. S2D–G), 2D iSPy Gaussian mixture with spherical covariance (yellow, (A–B) and Fig. 2C), and the 2D iSPy Gaussian mixture with diagonal covariance (blue, (C–E)). Uncertainty bars in all plots represent nuclei that may be classified incorrectly (log-likelihood probability less than 0.8) and nuclei that could be classified in another component (log-likelihood probability greater than 0.2) (see Methods).

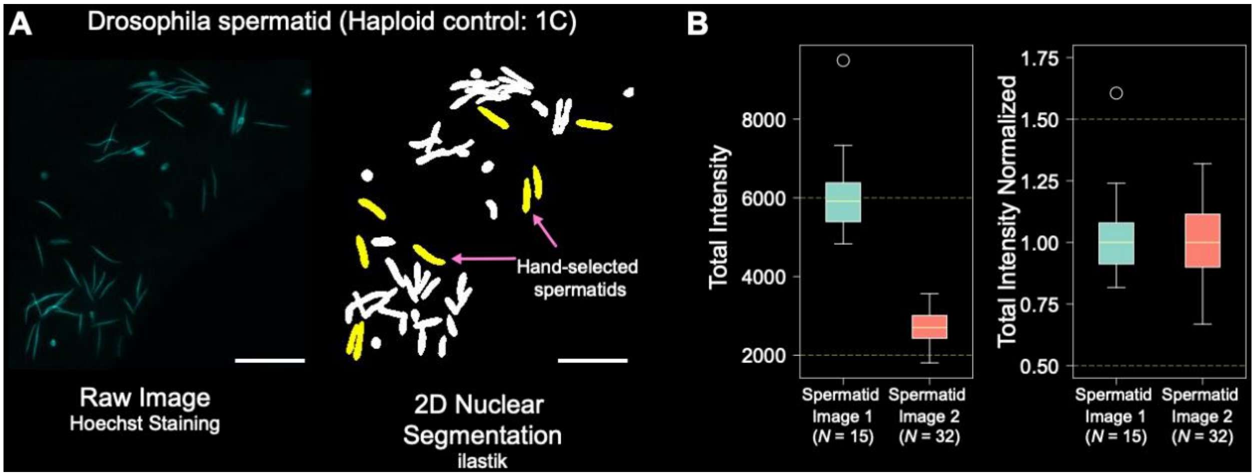


**Figure S6: Quantification of haploid sperm nuclei from Drosophila provides a control.** **(A)** Left: representative sum-projection of a confocal image of Drosophila spermatids stained with Hoechst. Right: segmented sperm nuclei using ilastik. Yellow nuclei signify hand-selected sperm which were complete and unobstructed (see Methods). Scale bars: 20 µm. **(B)** Example quantification of two spermatid images from the same experiment and slide. Left: total intensity of hand-selected spermatids from two different images. Both have sample sizes larger than $N = 10$ and medians between 2,000 and 6,000 (dashed green lines). Right: normalized total intensity of both images. All data points were divided by the median total intensity from each image. We mandate that at least 90% of normalized total intensity values are between 0.5 and 1.5 (dashed green lines). We found that 14/15 (93%) of the spermatids from the first image fit this criterion and 32/32 (100%) of spermatids from the second image fit this criterion. See Methods for a more detailed explanation.

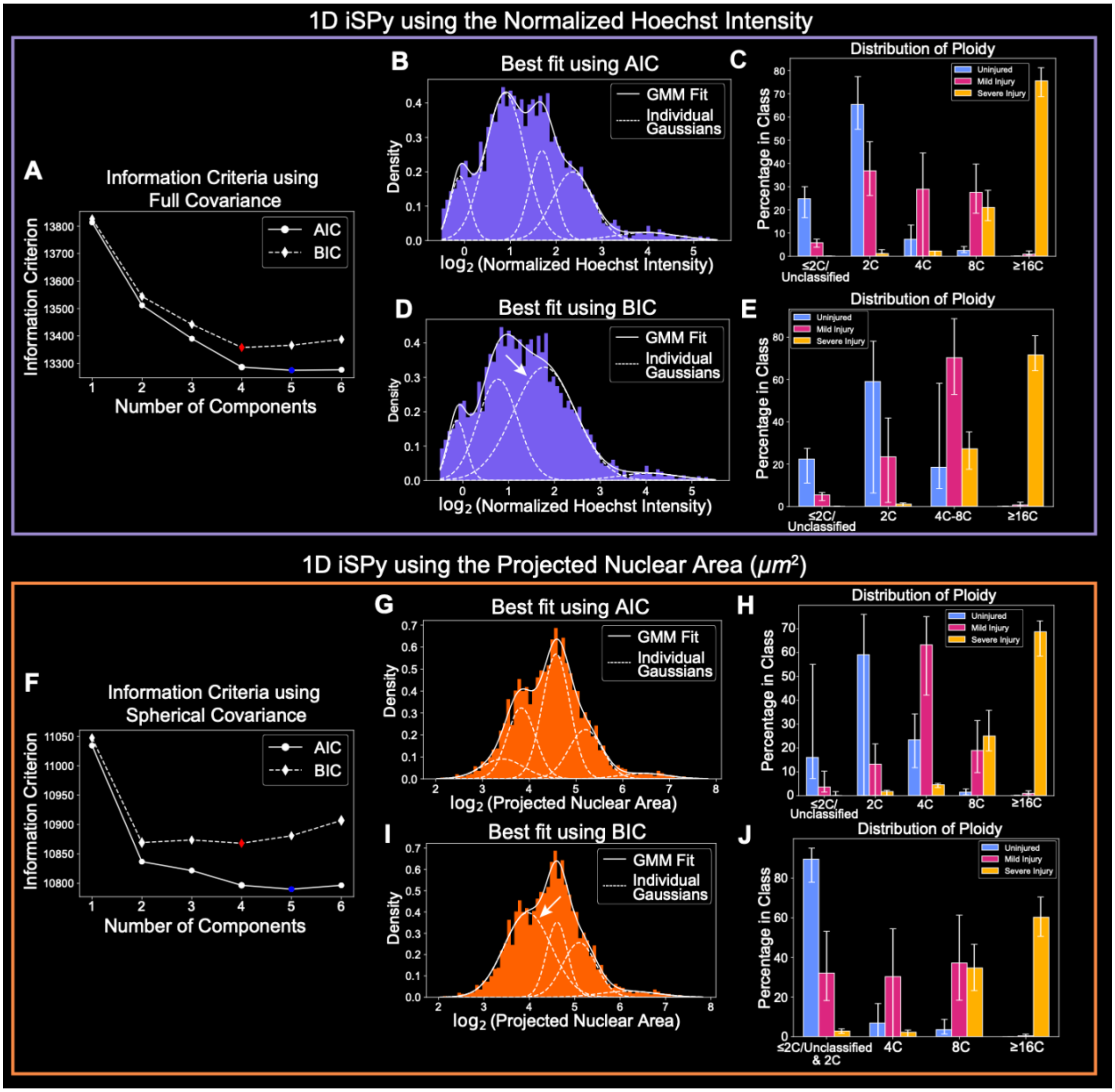


**Figure S7: AIC and BIC produce different best fits for Drosophila pyloric cells.** **(A–E)** A 1D Gaussian mixture for the normalized Hoechst intensity (see Methods). **(A)** Values of AIC (solid line) and BIC (dashed line) as the number of components is varied while using the full covariance matrices. The blue and red colored points represent the lowest AIC (five components) and BIC (four components), respectively. **(B)** The best fit of the 1D Gaussian mixture using AIC displaying the histogram (purple), the full Gaussian mixture fit (solid white line), and the five components (dashed lines). **(C)** The proportion of uninjured pyloric cells (blue), mildly injured pyloric cells (red), and severely injured pyloric cells (blue) in each predicted ploidy class: ≤2C/Unclassified, 2C, 4C, 8C, and ≥16C. **(D)** The best fit of the 1D Gaussian mixture using BIC displaying the histogram (purple), the full Gaussian mixture fit (solid white line), and the four components (dashed lines). Note the large component (arrow) that marks cells between 4C and 8C. **(E)** The proportion of uninjured pyloric cells (blue), mildly injured pyloric cells (red), and severely injured pyloric cells (blue) in each predicted ploidy class: ≤2C/Unclassified, 2C, 4C–8C, and ≥16C. **(F–J)** A 1D

Gaussian mixture for the projected nuclear area (see Methods). **(F)** Values of AIC (solid line) and BIC (dashed line) as the number of components is varied while using the spherical covariance matrices. The blue and red colored points represent the lowest AIC (five components) and BIC (four components), respectively. **(G)** The best fit of the 1D Gaussian mixture using AIC displaying the histogram (orange), the full Gaussian mixture fit (solid white line), and the five components (dashed lines). **(H)** The proportion of uninjured pyloric cells (blue), mildly injured pyloric cells (red), and severely injured pyloric cells (blue) in each predicted ploidy class: ≤2C/Unclassified, 2C, 4C, 8C, and ≥16C. **(I)** The best fit of the 1D Gaussian mixture using BIC displaying the histogram (orange), the full Gaussian mixture fit (solid white line), and the four components (dashed lines). Note the large component (arrow) marks cells between ≤2C/Unclassified and 2C. **(J)** The proportion of uninjured pyloric cells (blue), mildly injured pyloric cells (red), and severely injured pyloric cells (blue) in each predicted ploidy class: ≤2C/Unclassified & 2C, 4C, 8C, and ≥16C. See Supplemental Data Set 1 for exact values for (C), (E), (H), and (J). All uncertainty bars represent nuclei that may be classified incorrectly (log-likelihood probability less than 0.8) and nuclei that could be classified in another component (log-likelihood probability greater than 0.2) (see Methods).

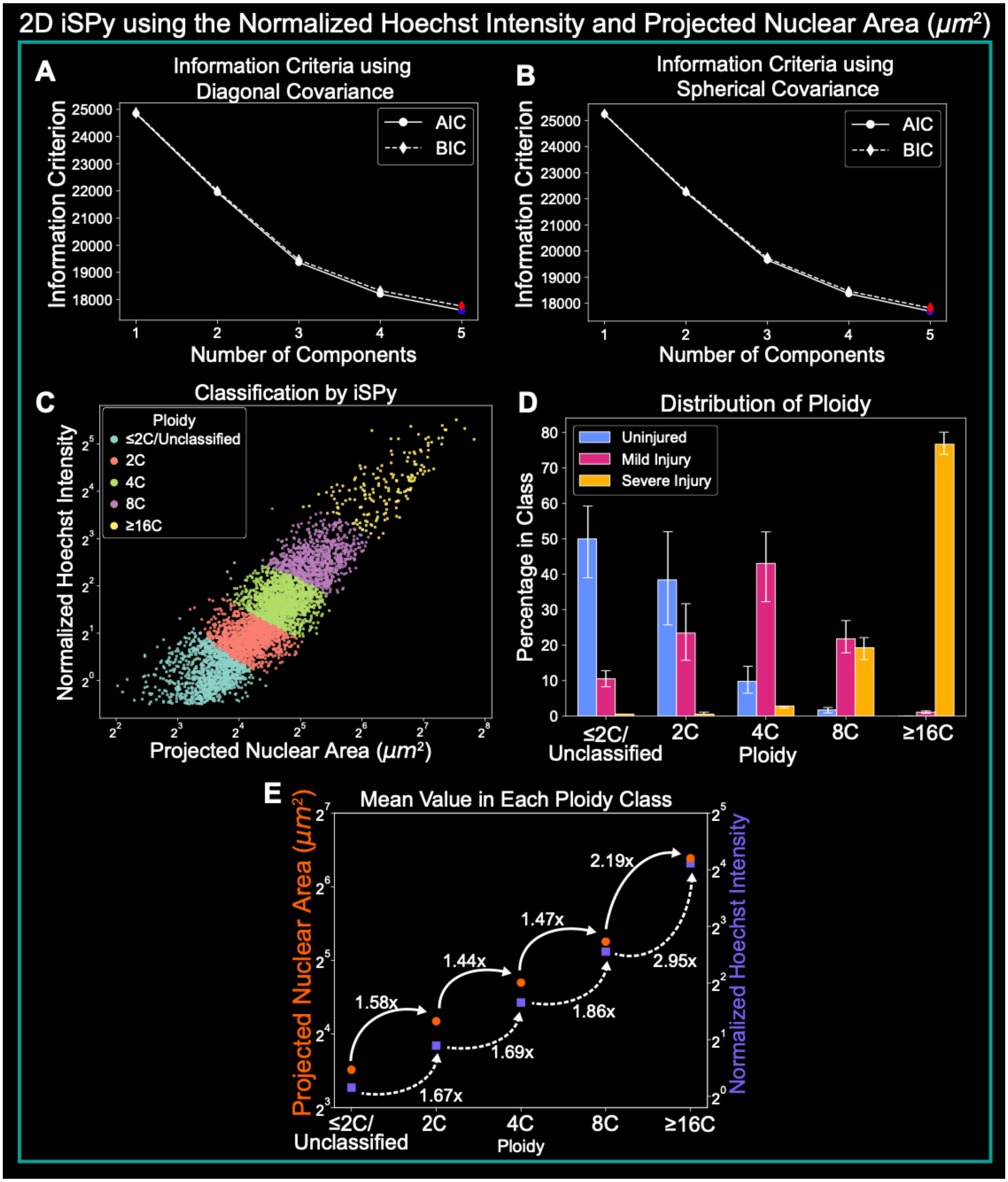


**Figure S8: The 2D Gaussian mixture for Drosophila pyloric cells predicts five components using either spherical or diagonal covariance matrices.** **(A–B)** Values of AIC (solid line) and BIC (dashed line) as the number of components is varied while using the **(A)** diagonal or **(B)** spherical covariance matrices. The blue and red colored points at five components represent the lowest AIC and BIC, respectively, and therefore the best fit. The best fit for (A) is shown in Fig. 3C. **(C)** iSPy prediction for each pyloric nucleus using the 2D Gaussian mixture with five components and spherical covariance matrix corresponding to ploidies of ≤2C/Unclassified, 2C, 4C, 8C, and ≥16C. **(D)** The proportion of pyloric cells in each ploidy class predicted from the 2D Gaussian mixture in (C) by severity of injury (uninjured, blue; mild injury, red; severe injury, yellow). Uncertainty bars represent nuclei that may be classified incorrectly (log-likelihood probability less than 0.8) and nuclei that could be classified in another component (log-likelihood probability greater than 0.2) (see Methods). See Supplemental Data Set 1 for exact values. **(E)** The mean of the normalized Hoechst intensity (purple squares, left axis) and projected nuclear area (orange circles, right axis) with the fold increase to the next ploidy class using the 2D Gaussian mixture in (C).

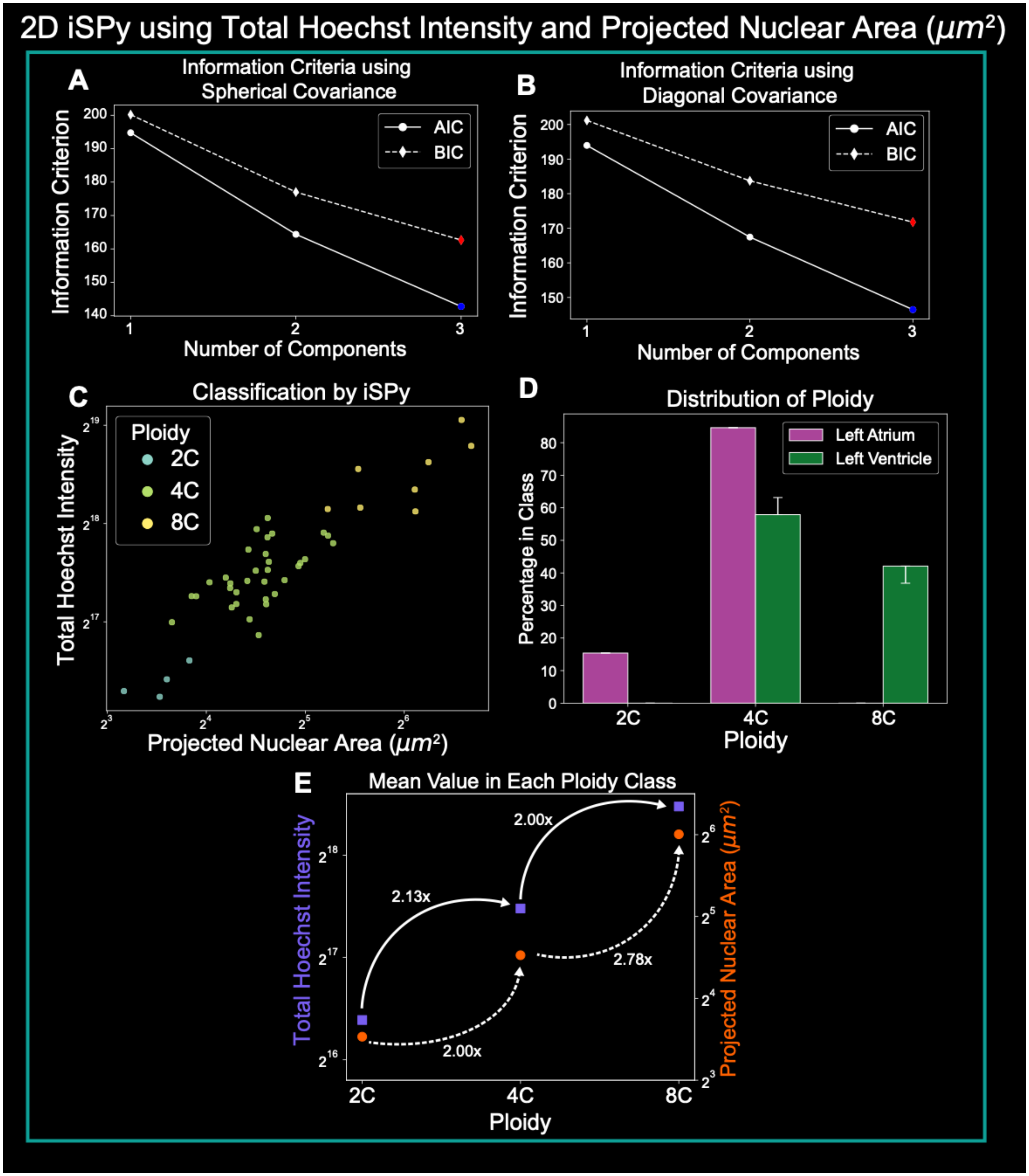


**<u>Figure S9: Using either spherical or diagonal covariance matrices predicts three ploidy classes for human cardiomyocytes.</u>** **(A–B)** Values of both information criteria AIC (solid line) and BIC (dashed line) as the number of components is varied while using the **(A)** spherical or **(B)** diagonal covariance matrices. The blue and red colored points at three components represent the lowest AIC and BIC, respectively, and therefore the best fit. The best fit for (A) is shown in Fig. 4C. **(C)** iSPy prediction for each pyloric nucleus using the 2D Gaussian mixture with three components and diagonal covariance matrix corresponding to ploidies of 2C, 4C, and 8C. **(D)** The proportion of cardiomyocytes in each ploidy class predicted from the

2D Gaussian mixture in (C) by heart chamber (left atrium, purple; left ventricle, green). See Supplemental Data Set 1 for exact values. Uncertainty bars represent nuclei that may be classified incorrectly (log-likelihood probability less than 0.8) and nuclei that could be classified in another component (log-likelihood probability greater than 0.2) (see Methods). **(E)** The mean of the Hoechst intensity (purple squares, left axis) and projected nuclear area (orange circles, right axis) with fold-increase to the next ploidy class using three components and diagonal covariance.
